# A scalable tumor-vasculature-on-chip for CAR T cell trafficking and efficacy studies

**DOI:** 10.64898/2026.02.05.703975

**Authors:** Luuk de Haan, Aleksandra Olczyk, Thomas Olivier, Joris Wesselius, Johnny Suijker, Claudia Al-Mardini, Todd Burton, Lenie van den Broek, Karla Queiroz

**Affiliations:** MIMETAS BV, De Limes 7, 2342DH Oegstgeest, The Netherlands; Leiden Academic Centre for Drug Research LACDR, Leiden University, The Netherlands

## Abstract

Most cellular therapies, like CAR T cells, remain ineffective in solid tumors. This is primarily due to a complex tumor microenvironment (TME), which creates biochemically hostile and often immunosuppressive conditions that limit efficacy of immunotherapies. Besides, cellular therapy efficacy is still often established in traditional 2D cultures that fail to simulate relevant aspects of solid tumor biology. Recent advances in three-dimensional (3D) and organ-on-chip culture systems have provided more physiologically relevant models for immunotherapy testing. These microphysiological systems (MPS) not only offer a 3D environment that alters tumor cell sensitivity to therapy but also enable inclusion of TME components and assessment of processes such as extravasation and infiltration, key steps in CAR T cell activity in vivo. This study focuses on applying an advanced culture technique and further building on the use of a scalable on-chip platform, the OrganoPlate, to grow EpCAM-positive and EpCAM-negative tumor cells in co-culture with an endothelial vessel to study EpCAM-targeting CAR T cell migration and killing kinetics. The CAR T cells specifically targeted and killed EpCAM-positive HT-29 tumor cells while EpCAM-negative A375 tumor cells were not affected. In addition, target cell killing was dependent on the ratio between CAR T and tumor cells (E:T ratio) and was enhanced by addition of IL-2. Inflammatory cytokines like INF-γ, TNFα and IL-6 increased overtime in cultures containing CAR T cells. Morphometric analyses of the endothelial compartment showed E:T ratio dependent disruption of endothelial vessels. Additionally, this system was able to distinguish EpCAM ScFv-CD28-CD3z and EpCAM ScFv-TM-4-1BB-CD3z CAR T cells killing abilities and was used for studying the effect of immune checkpoint inhibitors and Temozolomide, a DNA targeting drug, on CAR T cell performance. Altogether, this work adds to the available advanced culture techniques for immunotherapy developers by describing a model that is modular, scalable, and suitable for phenotypic and functional characterization of CAR T cells.

## 1. Introduction

Solid tumors remain one of the leading causes of death worldwide, and many of these diseases are deemed incurable or lack effective therapeutic options (1). Solid tumor tissues are complex entities composed of many tumor cell subpopulations as well as a diverse and dynamic cellular microenvironment that supports tumor cell growth and survival. A major breakthrough that is revolutionizing treatment of many types of cancer occurred with the development of immunotherapies, aimed at harnessing the immune system to treat cancer (2). Among these are checkpoint inhibitors, which are designed to prevent suppression of the immune infiltrate and thereby promote the anti-tumor response (3). However, many solid tumors exhibit low levels of immune cell infiltration or lack immune cells altogether and these so-called cold tumors usually show little response to checkpoint inhibition (4). One promising strategy to overcome this lack of immune infiltrate is adoptive cell therapy, particularly the use of Chimeric Antigen Receptor (CAR) T cells (5). These engineered T cells express synthetic receptors that redirect polyclonal T cells to specific surface antigens, enabling targeted tumor elimination (6). CAR T cell therapy has shown remarkable efficacy in treating relapsed and refractory B cell malignancies (7). However, despite these successes in treatment of hematological malignancies, the clinical efficacy of CAR T cells for solid tumors remains limited (8). Solid tumors present several physical and chemical barriers that are absent in hematological malignancies, like dysfunctional tumor vasculature and a potential hostile and immunosuppressive tumor microenvironment (TME) characterized by oxidative stress, nutritional depletion, acidic pH and hypoxia as well as the presence of inhibitory soluble factors, cytokines and suppressive immune cells (9). These cause a lack of CAR T cell trafficking into the tumor tissue as well as limited effector function and, together with antigen heterogeneity and on-target, off-tumor toxicity (OTOT), form major obstacles that affect CAR T cell efficacy in solid tumors (10).

Currently, several strategies are being developed and tested to enhance CAR T cell function, persistence, homing, and infiltration in the context of solid tumors. Traditional 2D in vitro models, despite their limitations, are still widely used in early-stage oncology drug discovery. In CAR T cell research, 2D cultures are commonly employed to assess efficacy through killing and cytokine release assays (11). However, these models fail to replicate the complexity of interactions between tumor cells and the TME as well as the surrounding healthy tissue, which collectively are known to influence immune cell extravasation, infiltration and function. As a result, 2D models often overestimate CAR T cell efficacy and provide limited insight into their behavior in 3D environments, properties that may correlate with CAR T cell clinical performance (12). Development of advanced *in vitro* models that allow testing of cellular therapies under more physiologically relevant conditions is therefore imperative. These models enable not only efficacy testing but also valuable readouts such as immune cell distribution and migration in 3D environments and will play a critical role in accelerating the preclinical development of cellular therapies by providing complex microenvironments that better mimic the biology of solid tumors. Organ-on-chip platforms have been leveraged to generate advanced tumor models that allow studying of immune cell recruitment as well as the testing of several immunotherapeutic modalities (13). Notably, on-chip systems have been used for CAR T cell efficacy and safety testing in the context of leukemia (Ma et al., 2025) and breast cancer (15). These studies are a great example of how such on-chip models can be used to advance our understanding of how cellular interactions and immunosuppressive factors influence CAR T cells in a clinical setting. However, they often lack the throughput needed for industrial application or require specialized equipment to establish (16).

Here, we describe an on-chip tumor model to evaluate CAR T cell migration and killing capacity using the OrganoPlate. We previously described a model comprising endothelium and T cells to study extravasation and migratory behavior of T cells (17). First, we adapted the above-mentioned model by incorporating extracellular matrix (ECM)-embedded colorectal cancer (CRC) cells. We then introduced commercially available EpCAM-targeting CAR T cells and assessed extravasation, infiltration and killing efficacy. The CAR T cells specifically targeted and killed EpCAM-positive HT-29 tumor cells while EpCAM-negative A375 tumor cells were not affected despite similar levels of CAR T cell infiltration. Clinically relevant inflammatory cytokines such as IFN-γ, TNFα and IL-6 increased overtime in cultures exposed to CAR T cells. We also showed that killing of the target cells is dependent on the ratio between the CAR T cells and the tumor cells and can be increased by IL-2 supplementation. Furthermore, morphometric analysis of the endothelial compartment showed dose-dependent disruption of the endothelial vessel. Additionally, the system was used to distinguish different CAR designs in functional performance.

Finally, the model setup was used to investigate the effects of combinations of immune checkpoint inhibitors and the DNA targeting agent Temozolomide, showing distinct responses depending on treatment and tumor cell type. Altogether, this study adds to the available advanced culture techniques for development of novel immunotherapies by providing scientists with a microphysiological system (MPS) that is modular, scalable, cost-effective and suitable for phenotypic and functional characterization of CAR T cells.

## 2. Material & Methods Cell culture

Human melanoma cell line A375-GFP (Innoprot, P20122) and colorectal cancer cell line HT-29-GFP (Innoprot, P20123) were cultured in T-75 flasks using DMEM (Sigma, D6546) supplemented with 10% Fetal Bovine Serum (Gibco, A5670801), 2 mM L-Glutamine (Sigma, G7513) and 1% Penicillin-Streptomycin (Sigma, P4333). Medium was replaced 24 hours after thawing with fresh medium containing 250 µg/mL G418 (Invivogen, ant-gn-1). Human colorectal cancer cell line HCT116-GFP (GeneCopoeia, SL024) was cultured in RPMI-1640 (Gibco, 11875093) supplemented with 10% Fetal Bovine Serum and 1 % Penicillin-Streptomycin. Medium was replaced 24 hours after thawing with fresh medium containing 1 µg/mL puromycin (Invivogen, ant-pr-1). Before seeding, tumor cell lines were harvested using 0.025% trypsin/EDTA (Lonza, CC-5012), counted using Trypan Blue (Thermo Fisher Scientific, 15-250-061) exclusion, pelleted (200g, 5 min) and resuspended in AIM-V medium (Gibco, 12055091). All cell culture was performed in a humidified incubator at 37 °C and 5% CO_2_.

Human umbilical vein endothelial cells (HUVEC, Lonza, C2519AS) were cultured in T-75 flasks using complete EGM-2 medium (Lonza, CC-3162). HUVECs were harvested at passage 5 as described above except they were resuspended in EGM-2 medium at a density of 1e7 cells/mL.

Human EpCAM scFv-TM-CD28-CD3ζ CAR T cells (ProMab, PM-CAR1020), human EpCAM-scFv-TM-4-1-BB- CD3ζ CAR T cells (ProMab, PM-CAR1068) and control T cells from healthy donor (ProMab, PM-CAR2003) were cultured in 6-well plates. EpCAM scFv-TM-CD28-CD3ζ were expanded using CAR T cell expansion medium (ProMab, PM-2000) according to supplier’s protocol. Surplus of CAR T cells was resuspended in CELLBANKER Cell Freezing Media (Amsbio, 11888) at 1e7 cells/mL, frozen and stored at -150 °C. EpCAM-scFv-TM-4-1-BB- CD3ζ could not be expanded. Prior to an experiment, CAR T cells and/or control T cells were thawed in 6-well plates and recovered overnight in AIM-V medium before use in the OrganoPlate.

### Generation of tumor-on-chip cultures

Tumor-on-chip cultures were generated using the MI-OR-CU OrganoReady^®^ Lumenised Collagen 3-lane 40 (MIMETAS, 0001L). The microfluidic chip consists of three different channels, of which the middle channel contains an ECM gel that is hollow (Fig. 1A, B and C). The model is composed of two distinct compartments: an endothelial compartment in the top channel of the microfluidic chip adjacent to a tumor compartment in the middle channel. The third channel does not contain cells and can be utilized to introduce chemotactic triggers.

**Figure 1.**
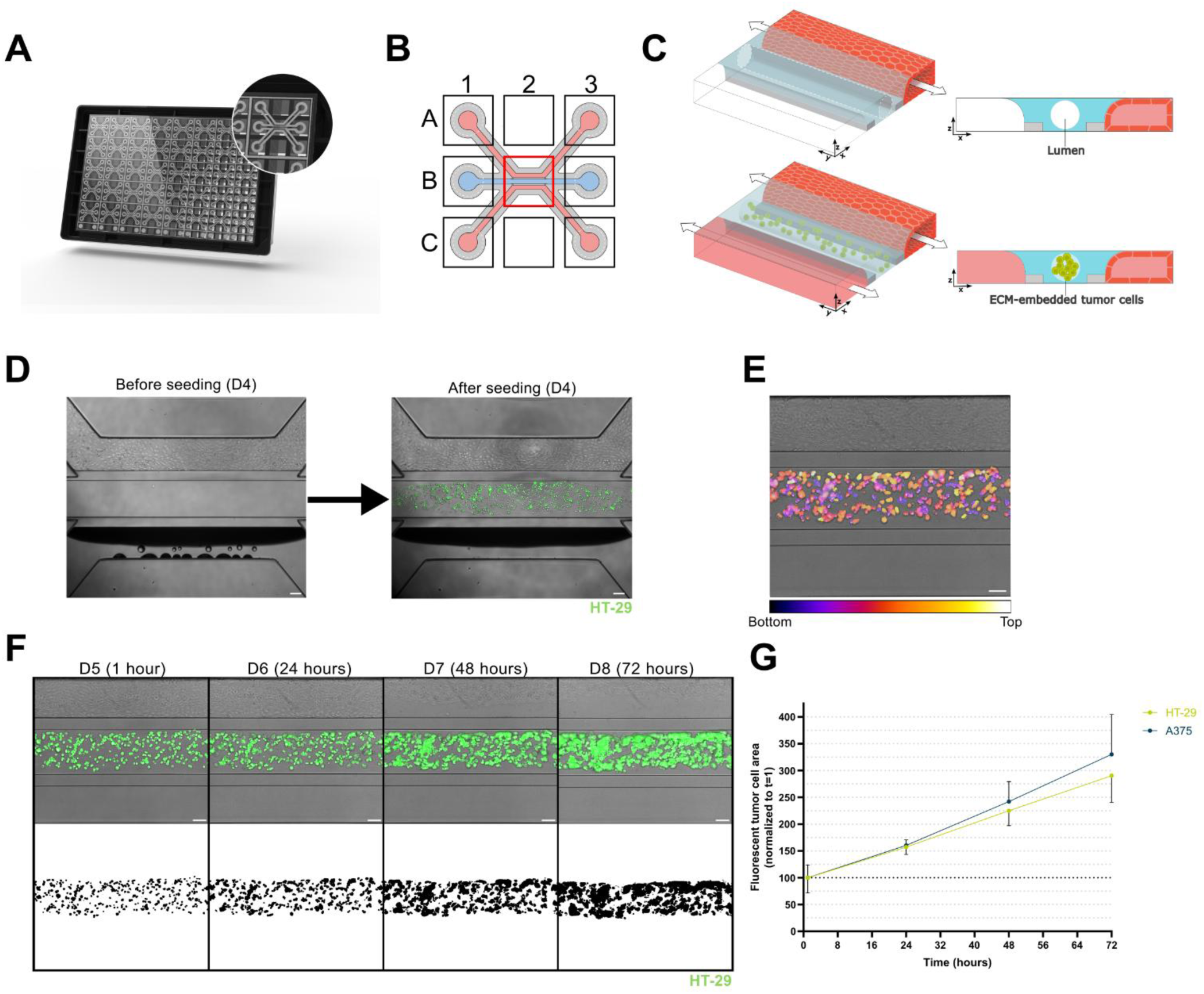
Development of an on-chip model comprising endothelium and ECM-embedded tumor cells. (A) OrganoPlate 3-lane 40, containing 40 individual microfluidic chips (zoom). (B) Schematic representation of a single microfluidic chip. The chip is made up of a 3 by 3 grid and consists of three microfluidic channels that come together in the center well (B2). The blue channel contains the lumenised extracellular matrix (ECM) gel and the red channels are used to either introduce endothelial cells or culture medium. (C) Schematic representation of the culture set-up. Endothelial cells are introduced in the top channel of the microfluidic chip and form a tubular structure against the lumenised ECM gel that resides in the middle channel. The lumen is indicated with the dashed lines. After complete formation of the endothelial vessel, tumor cells, embedded into a second ECM gel, are introduced in the lumen. (D) Images of an endothelial vessel grown against the lumenised ECM for 4 days, before (left) and after (right) introduction of HT-29-GFP tumor cells. (E) HT-29-GFP tumor cells are distributed in 3D after introduction into the lumenised ECM. Z-planes are color coded. (F) HT-29-GFP tumor cells proliferate after introduction in the lumenised ECM and cause regression of the endothelial vessel over time. The fluorescent signal could be used to create a binary mask that was used to track growth of HT-29-GFP tumor cells from day 5 until day 8. (G) Quantification of tumor cell growth based on binary masks. Fluorescent area at each timepoint was normalized against the area at t=1 for each individual replicate (HT-29: N=8, n=27, A375: N=2, n=8). Data shown in the graphs are presented as mean ± SD. Scale bars indicate 100 µm.

HUVECs, harvested as described above, are introduced in the top perfusion channels by loading 2 µL cell suspension at a density of 1e7 cells/mL into the inlet, followed by 50 µL of EGM-2. The OrganoPlate is placed in the MIMETAS plate stand for 2 hours to promote attachment of the cells to the ECM gel. Upon attachment, 50 µL of EGM-2 medium is added to the outlet. The OrganoPlate is then placed on an OrganoFlow^®^ rocking platform (MIMETAS, MI-OFPR-L) at 7 degrees inclination with an 8-minute interval to induce perfusion.

On day 4, when endothelial vessels were fully developed, tumor cells were harvested and embedded in an ECM at a density of 5.7e6 cells/mL. ECM was prepared by mixing rat tail collagen I (5 mg/mL, Cultrex, 3447-020-01), HEPES (1M, Thermo Fisher Scientific, 15630056) and NaHCO_3_ (37 g/L, Sigma, S5761) in a ratio of 8:1:1 and addition of 10% Fibronectin (1 mg/mL, Sigma, F0895), resulting in a mix with a final concentration of 3.6 mg/mL collagen I + 0.1 mg/mL fibronectin. After washing the middle channel with PBS, tumor cells were loaded in the lumenised compartment in the middle channel of the microfluidic chip by placing 30 µL of AIM-V medium on the inlet followed by 0.7 µL of ECM mix on the outlet, resulting in the introduction of 4.000 tumor cells per microfluidic chip. The plate was placed in a humidified incubator (37 °C and 5% CO_2_) for 12 minutes to polymerize the introduced ECM gel. Finally, 50 µL of AIM-V medium was added to inlet (top-up) and outlet of the middle channel before the plate was placed back on the rocker. Co-cultures were grown overnight before addition of CAR T cells or control T cells.

### Flow cytometry

Composition of stored CAR T cells was assessed after thawing and recovery. CAR T cells were transferred to a 96-well V-bottom plate and spun down (300g, 5 min). Samples were stained with Zombie NIR™ Fixable Viability Kit (Biolegend, 423106) for 30 minutes at room temperature (RT) in the dark. Samples were blocked with FcR blocking reagent Human (Miltenyi Biotec, 130-059-901) and stained for 30 min at 4°C in the dark. Antibodies used include BUV605 anti-CD3 (Biolegend, clone UCHT1, 300460), BUV395 anti-CD4 (BD Biosciences, clone SK3, 563550) and BV785 anti-CD8 (Biolegend, clone SK1, 34470). After staining, samples were spun down (300g, 5 min) and incubated with eBioscience™ Foxp3/Transcription Factor Staining Buffer Set (Invitrogen, 00-5523-00) for 45 minutes at 4°C in the dark. Finally, samples were stained intracellularly with BV421 anti-Ki-67 (BD Biosciences, clone B56, 562899) for 30 minutes at RT in the dark. Samples were washed and kept in FACS buffer at 4°C in the dark until analysis within 24 hours. UltraComp eBeads™ Plus Compensation Beads (Invitrogen, 01-3333-42) were stained separately with single antibodies and used as reference controls to determine antibody spectral signatures. Cells stained with Zombie NIR™ Fixable Viability Kit were used to set spectral signature of detect dead cells. Data acquisition was acquired on a Cytek Aurora 3-laser 16V-14B-8R spectral flow cytometer using Cytek Bioscience SpectroFlo software. Signal unmixing was performed automatically by the flow cytometer and manually adjusted. Analysis was performed using FlowJo 10 (FlowJo LLC, v10.10.0).

### Addition of CAR T cells to tumor-on-chip cultures

The endothelial compartment was exposed to 4 ng/mL TNFα (Immunotools, 11343015) in EGM-2 medium for 4 hours. Meanwhile, CAR T cells were collected from 6-well plates and counted using a C-chip disposable hemocytometer (NanoEntek, DHC-N01). In case of tracking experiments, CAR T cells were spun down (300g, 5 min) and resuspended in AIM-V containing freshly prepared 0.5 µM DiR (Invitrogen, D12731). After vortexing, cell suspension was incubated in the dark at 37 °C for 20 minutes before staining was terminated by addition of an equal volume of AIM-V medium containing 10% serum. Suspensions were spun down (300g, 5 min) and CAR T cells were resuspended at a density of 5.7e5 cells/mL from which different working concentrations (E:T ratios) were prepared depending on the experiment. After 4 hours of exposure, medium containing TNFα was removed and 70 µL AIM-V containing CAR T cells was added to the inlet of the top perfusion channel, followed by addition of 70 µL AIM-V to the outlet. In case of IL-2 supplementation, AIM-V containing 100 IU/mL was added to the outlet, ultimately resulting in 50 IU/mL IL-2 in the chip after mixing. In case of experiments with checkpoint inhibitors and Temozolomide, AIM-V containing 20 µg/mL Nivolumab (MedChemExpress, HY-P9903) and/or Ipilimumab (Selleckchem, A2001) and/or 150 µM Temozolomide (MedChemExpress, HY-17364) was added to the outlet, ultimately resulting in 10 µg/mL and/or 75 µM in the chip after mixing. Finally, medium was aspirated from the middle channel and 70 µL of either plain AIM-V or AIM-V containing 800 ng/mL CXCL12 (Peprotech, 300-28A) was added to the inlet and outlet of the bottom perfusion channel. Optionally, Draq7 (1:2000, Biostatus, DR71000) was added into the culture medium as an indicator of dead cells.

### CAR T cell infiltration and killing assay

After apically introducing CAR T cells to the endothelial vessel, their migration behavior and/or killing abilities were tracked over time (24-72h). Several CAR T and tumor cell ratios were tested (1:2, 1:1, 2:1, 5:1 and 10:1). Images of the live tumor-on-chip co-cultures were acquired using the 60 µm spinning disc confocal mode, a 0.45NA dry-air 10× objective and dichroic/emission fluorescent filters for FITC, Cy5 and Cy7. Maximum intensity projections of fluorescently labelled CAR T cells were generated and used to determine positions (top to bottom) of the CAR T cells in the microfluidic chips using FIJI (version 2 build 1.52d). Images were corrected for photobleaching-induced artifacts, and a rolling ball background correction was applied to improve the signal-to-noise ratio. An automatic thresholding routine was used to create binary masks of fluorescently labelled CAR T cells within images, and a particle detection algorithm was applied resulting in individual CAR T cells being outlined and labelled. The list was used to extract X/Y positional information about individual CAR T cells within microfluidic chips. CAR T cells quantified at Y positions ≤ 450 µm were considered to reside in endothelial vessels, while CAR T cells quantified at Y positions > 450 µm were considered migrated CAR T cells. Maximum intensity projections of GFP-positive tumor cells were generated, and binary masks were created using thresholding (Otsu’s method) based on control chips (no CAR T cells). In case of HCT116 tumor cells, Contrast Limited Adaptive Histogram Equalization was used prior to generating the binary masks. Masks were used to extract total tumor area at different timepoints, and growth of tumor cells was assessed by correcting against the area at t=1.

### Immunocytochemistry

At endpoint, tumor-on-chip cultures were fixed as follows; all medium was removed from each well, replaced with fixation solution 3.7% formaldehyde (Sigma, 252549) in HBSS with calcium and magnesium (Gibco, 14025092), and incubated at RT for 15 minutes. Fixation solution was removed and each well was washed three times with phosphate buffered saline (PBS). Following the third wash, 50 µL PBS was added to each well and the plate was sealed using Parafilm and stored at 4°C until immunofluorescent staining. Plates were permeabilized and blocked for 2 hours with blocking buffer (1% Triton X-100 + 3% BSA in PBS), followed by overnight incubation with primary antibodies in antibody buffer (0,3 % Triton X-100 + 3% BSA in PBS). Cultures were washed three times with washing buffer (0.3% TritonX-100 in PBS), followed by 2-hour incubation with secondary antibodies and Hoechst (1:2000, Invitrogen, H3570) in antibody buffer. After washing three times with washing buffer and once with PBS, PBS was added to all wells and the plate was imaged using the ImageXPress XLS Micro Confocal (Molecular Devices). Primary antibodies used: VE-cadherin (1:1000, Abcam, Ab33168) and CD45 (1:100, R&D systems, MAB1430). Secondary antibodies used: Goat anti-Rabbit Alexa Fluor 555 (1:250, Invitrogen, A-21428) and Goat anti-Mouse Alexa Fluor 750 (1:250, Invitrogen, A-21037). Maximum intensity projections were generated and saved for further analysis. Bottom and top layers of the endothelial vessel were acquired separately for VE-cadherin morphometric analysis.

### Morphometric analysis of VE-cadherin network

IN Carta® (v2.6.01082563) was used to develop an image analysis pipeline. A Flexi-Protocol 2D pipeline was designed that analyses the different aspects of the endpoint immunofluorescent staining data. Nuclei, and individual cell-areas based on VE-cadherin were segmented using custom deep-learning models through the SINAP module in IN Carta. Touching objects were split using a watershed algorithm. These models are based on a U-Net CNN architecture with a Resnet backbone (18–20).

VE-cadherin objects were described using a range of different morphological and spatial descriptors (Supplementary Table 1). Morphological parameters included: Area, Gyration Radius, Compactness, Form Factor, Elongation, Chord Ratio, Diameter, and Perimeter. Spatial parameters included were: Minor Axis, Major Axis, Major Axis Angle. Additionally, the number of intact cells were captured in an object count labelled “Count”. The ratio between the total area covered by cells, and the total available area in the channel was represented as Confluence.

The VE-cadherin objects were cleaned up during pre-processing to reduce the impact of artifacts on the analysis. Cells had to be between 100µm and 2000µm to be considered valid, and a 95^th^ percentile cut-off on the above listed measurements was used to filter out outliers.

Systemic tubular degradation is captured by the Count measurement, as a decrease in mean area of individual cells or as a change in confluency. Tubular reorganization, or a change in alignment against the flow, is captured by shape-changes (Form Factor, Compactness, Elongation, Major Axis Angle). Jointly these measurements describe a more localized “shrinking” of cells versus the complete loss of cell segments from the tubule.

### Quantification of cytokine secretion

Cytokine profiles of medium samples taken from co-cultures were assessed using a 6-plex Human Discovery Assay (R&D Systems, LXSAHM-06). The samples were diluted 5x or 2x depending on number of administered CAR T cells before being analyzed for levels of IFN-γ, IL-2, IL-6, TNFα, soluble PD-L1/B7-H1 and Granzyme B according to the protocol provided by the manufacturer using Luminex MAGPIX and xPONENT software. Raw data was processed using Quantist software (R&D Systems, QUANTIST).

### Principal Component Analysis (PCA) and multi-parameter data representation

Data of Granzyme B, IL-6, soluble PD-L1/B7-H1 and TNFα levels from the Luminex analysis were first normalized to the CAR T cell control, thereby represented as a fold-change compared to untreated co-cultures. IL-2 and IFN-γ levels were too low to be quantified accurately and therefore omitted from the analysis. Similarly, endpoint tumor area measurements based on GFP expression were normalized to the CAR T cell control. Python (v3.10.6), with the packages numpy (v1.26.4), seaborn (v0.13.2) and scikit-learn (v1.6.1), was used to perform the Principal Component Analysis (PCA) on this collective dataset. A biplot was made using plotly-express (v5.24.1) to visualize the direction and contribution of the different features in the PCA.

To assess treatment effects across multiple parameters, pairwise distances were computed and visualized as a matrix using Python (v3.10.6), with the packages: numpy (v1.26.4), seaborn (v0.13.2), plotly (v5.24.1), sci-kit learn (v1.6.1), scipy (v1.13.1). Unsquared correlation distance was chosen as it emphasizes feature patterns rather than magnitude of the effect.

### Additional data analysis

General data handling was performed in Excel and graphs were generated in GraphPad Prism 10. Statistics are conducted in GraphPad Prism. Biological and/or technical replicates are listed in figure legends.

## 3. Results

### Development of a modular tumor-vasculature microphysiological system

Endothelial vessels were grown in the OrganoPlate 3-lane 40, which contains 40 individual microfluidic chips (Fig. 1A and B), by introducing HUVEC cells in the top channel of the chip and culturing them against a lumenised ECM hydrogel under perfusion flow (Fig. 1C). After 4 days of culture, the endothelial vessel was fully formed and ECM-embedded HT-29-GFP tumor cells, a colorectal adenocarcinoma cell line, were successfully introduced into the lumen without disrupting the endothelial vessel (Fig. 1C and D). Utilizing the GFP expression by the HT-29 cells, the distribution of the tumor cells in the middle channels of the microfluidic chips was assessed. Confocal microscopy revealed the HT-29-GFP tumor cells to be present in various planes, indicating three-dimensionality of the culture (Fig. 1E). Variability of the tumor cell seeding was assessed by calculating the tumor cell coverage of the middle channel, indicating an average Area% of 23.41±7.49 based on 148 microfluidic chips across 4 different OrganoPlates (Sup. Fig. 1A). Based on the GFP expression, a semi-automated quantification method to track HT-29-GFP tumor cell growth was established (Fig. 1F). Starting day 5, the co-culture was followed for 72 hours, showing a 2.9-fold increase in HT-29-GFP tumor cell area (Fig. 1G, green line). The HT-29-GFP tumor cells could easily be substituted with A375-GFP melanoma cells (Sup. Fig. 1B), resulting in a similar increase in A375-GFP tumor cell area over 72 hours of culture (Fig. 1G, blue line). Interestingly, in case of both HT-29-GFP and A375-GFP tumor cells, the endothelial vessel was seen to regress after 48 hours of co-culture (Fig. 1F and Sup. Fig. 1B). This 72-hour co-culture period, during which the tumor cells grow and the endothelial vessel regresses, provides an experimental window that can be used for co-culture with CAR T cells.

### CAR T cell trafficking towards ECM-embedded tumor cells

According to the Human Protein Atlas, HT-29-GFP tumor cells express epithelial cell adhesion molecule (EpCAM) whereas the melanoma cell line A375-GFP does not (22,23). Commercially available CAR T cells targeting EpCAM, from here onwards called EpCAM-CD28 CAR T cells, contain a roughly equal distribution of CD4⁺ and CD8⁺ T cell subsets (Fig. 2A and Sup. Fig. 2). After being perfused through the endothelial vessel, CAR T cells adhered to the vessel wall and underwent transendothelial migration, ultimately extravasating and migrating towards the tumor compartment (Fig. 2B, C and D). EpCAM-CD28 CAR T cells were fluorescently labelled before being apically added to the endothelial vessels to track extravasation and infiltration into the tumor compartment (Fig. 2E, F and G). EpCAM-CD28 CAR T cells were found to infiltrate both HT-29-GFP and A375-GFP tumor compartments over the course of the experiment in low numbers. However, just as it was found in previous research using healthy T cells (de Haan et al., 2021), migration of EpCAM-CD28 CAR T cells increased significantly in the presence of a gradient of CXCL12, which was introduced into the system via the channel opposite of the endothelial vessel. While EpCAM-CD28 CAR T cells infiltrated both HT-29-GFP and A375-GFP tumor compartments, a difference in migratory behavior was seen between the two on-chip models. The total number of EpCAM-CD28 CAR T cells showed a stronger increase over time in the presence of A375-GFP tumor cells (Fig. 2F) despite similar numbers at the start of the co-culture. However, the percentage of EpCAM-CD28 CAR T cells that infiltrated the tumor compartment continued to increase overtime for HT-29 cultures, while infiltration percentage seemed to reach a plateau after 24 hours in case of A375 cells (Fig. 2G).

**Figure 2.**
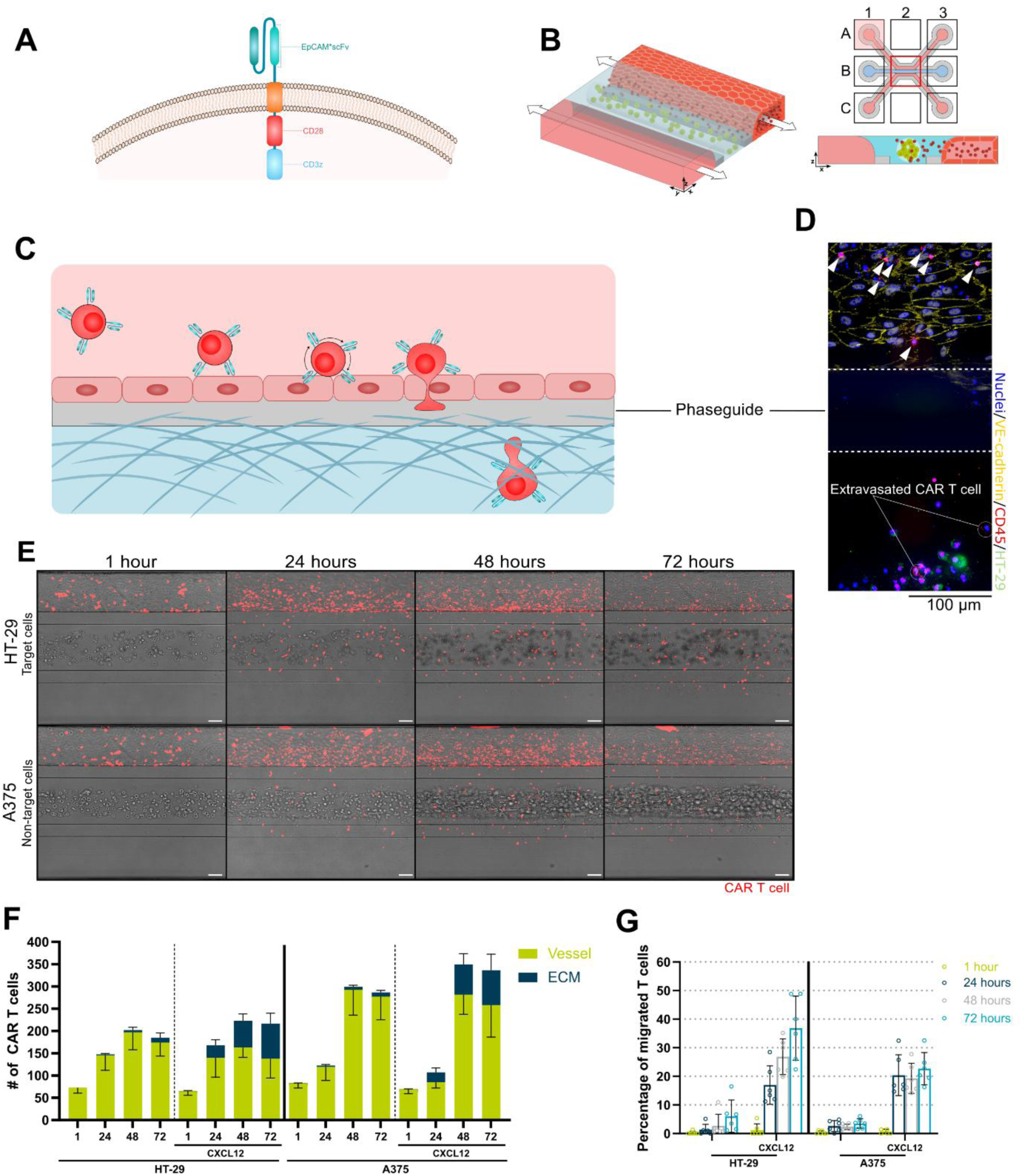
Migration of EpCAM-CD28 CAR T cells towards HT-29 and A375 tumor compartments. (A) Schematic representation of EpCAM-CD28 CAR. The CAR construct consists of an antigen-binding domain (EpCAM*scFv) fused to a transmembrane domain (orange) and two intracellular signaling domains (CD28, red and CD3z, blue). (B) Schematic representation of the co-culture set-up. EpCAM-CD28 CAR T cells (red) are perfused through the endothelial vessel via addition to the perfusion inlet (A1). Upon addition, CAR T cells adhere to the endothelial vessel wall and infiltrate the tumor compartment after undergoing the process of transendothelial migration. (C) Transendothelial migration process. CAR T cells (red) adhere to the endothelial vessel wall and, after rolling along the layer of endothelial cells, ultimately extravasate and infiltrate the adjacent ECM gel (blue = ECM fibrils). (D) Maximum intensity projection of cultures stained for VE-cadherin (yellow) and CD45 (red). Arrowheads indicate CAR T cells that are residing in the endothelial vessel. Circled CAR T cells have extravasated and reside close to HT-29-GFP cells (green). (E) Migration of fluorescently labelled EpCAM-CD28 CAR T cells towards HT-29 and A375 tumor compartments in the presence of a CXCL12 gradient over 72 hours of co-culture. (F) Quantification of the number of fluorescently labelled EpCAM-CD28 CAR T cells during 72 hours of co-culture with either HT-29 or A375 tumor cells (n=6). (G) Percentage of migrated EpCAM-CD28 CAR T cells during 72 hours of co-culture with either HT-29 or A375 tumor cells. Percentage is calculated as ‘# of CAR T cells in ECM/# of CAR T cells in Vessel + ECM’ (n=6). Data shown in the graphs are presented as mean ± SD. Scale bars indicate 100 µm.

### EpCAM-CD28 CAR T cells specifically recognize and kill target cells

Recognition of the target antigen by extravasated CAR T cells triggers the release of effector molecules like Granzyme B and IFN-y, leading to apoptosis of the targeted cell (Fig. 3A). Tumor growth was quantified using the GFP-based method that was described before for both A375-GFP (Fig. 3B and C) and HT-29-GFP (Fig. 3B and D) tumor-on-chip models. Addition of EpCAM-CD28 CAR T cells resulted in a slight decrease of 18.6% in A375-GFP tumor cell growth after 72 hours compared to control (Control: 305.63±82.88 vs. CAR T cells: 248.66±40.12). A similar 16.7% decrease in A375-GFP tumor cell growth was observed in the case of untransduced T cells (T cells: 254.47±56.79). In contrast, HT-29-GFP tumor cell growth was significantly impaired after the addition of EpCAM-CD28 CAR T cells, resulting in a 95.6% decrease compared to control after 72 hours of co-culture (Control: 262.94±34.81 vs. CAR T cells: 11.46±10.58). However, the addition of untransduced T cells only caused a minor decrease of 5.7% (T cells: 247.9±37.63).

**Figure 3.**
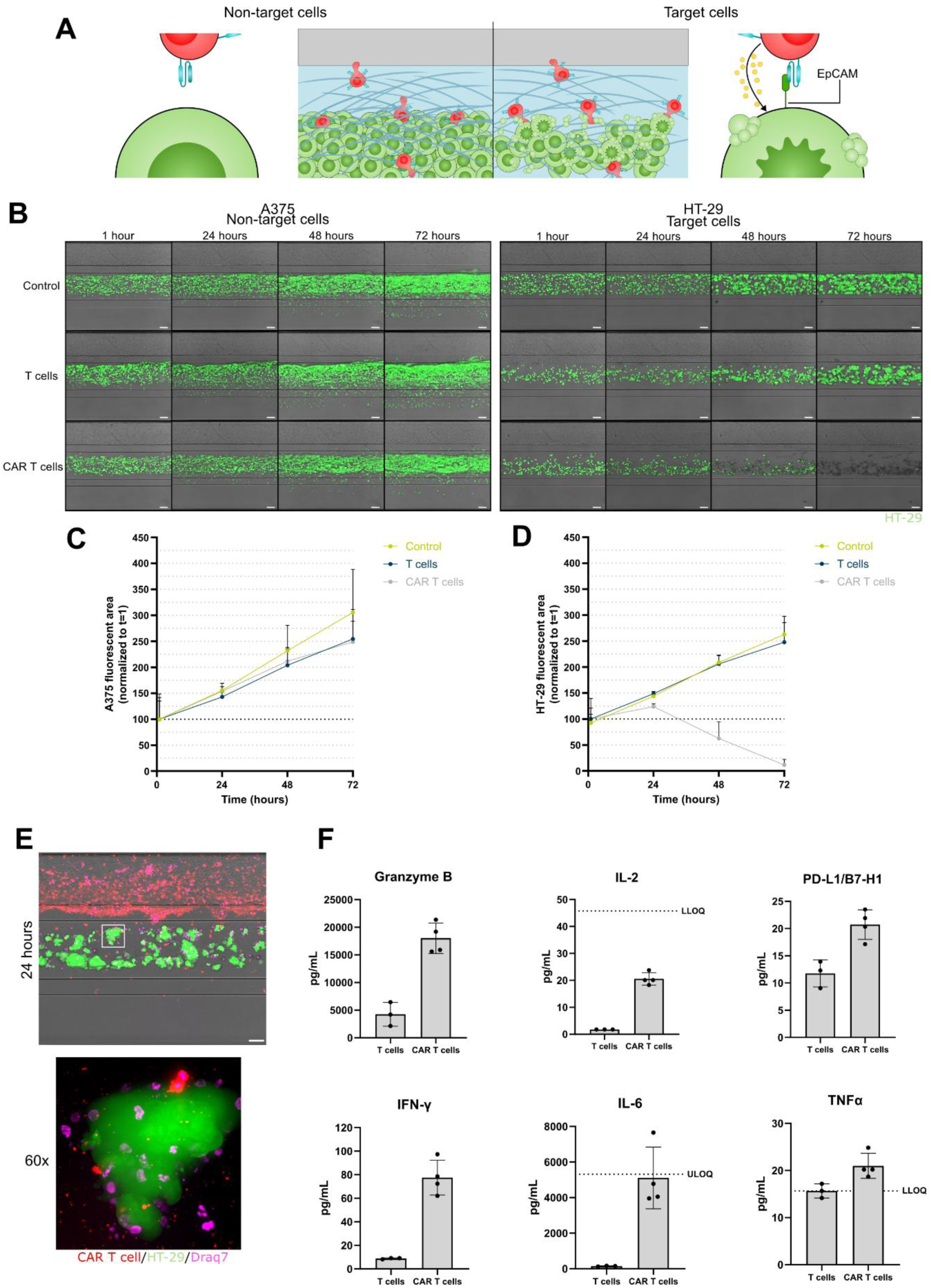
EpCAM-CD28 CAR T cells cause specific killing of HT-29-GFP tumor cells and secretion of inflammatory mediators. (A) Schematic representation of CAR T-mediated target cell killing. Upon extravasation, EpCAM-CD28 CAR T (red) cells infiltrate the adjacent ECM and encounter tumor cells (green). EpCAM-positive tumor cells will be recognized by the CAR T cells, resulting in the release of effector molecules and the killing of the targeted cell. (B) Images of A375-GFP and HT-29-GFP tumor cells co-cultured with untransduced T cells or EpCAM-CD28 CAR T cells for 72 hours. (C) Quantification of A375-GFP tumor cell growth during co-culture with either untransduced T cells (blue line) or CAR T cells (grey line). Fluorescent area at each timepoint was normalized against the area at t=1 for each individual replicate (n=4). (D) Quantification of HT-29-GFP tumor cell growth during co-culture with either untransduced T cells (blue line) or CAR T cells (grey line). Fluorescent area at each timepoint was normalized against the area at t=1 for each individual replicate (n=4-6). (E) Maximum intensity projection of HT-29-GFP tumor cells (green) after 24 hours of co-culture with EpCAM-CD28 CAR T cells (red). Dead cells are identified by Draq7 signal (magenta). White box depicts region that was assessed with confocal microscopy at 60x magnification (bottom panel). (F) Analysis of cytokine levels (pg/mL) in media samples after 72 hours of co-culture of HT-29-GFP tumor cells with either untransduced T cells (n=3) or EpCAM-CD28 CAR T cells (n=4). Data points outside of lower (LLOQ) and upper (ULOQ) limits of detection (dashed lines) are extrapolated from the standard curve. Data shown in the graphs are presented as mean ± SD. Scale bars indicate 100 µm.

Fluorescently labeled EpCAM-CD28 CAR T cells can be visualized in both the endothelial and tumor compartment and can be seen interacting with HT-29-GFP clusters within the tumor compartment (Fig. 3E). Analysis of culture media from HT-29-GFP cultures after the 72-hour co-culture showed strongly increased levels of effector molecules Granzyme B and IFN-γ, both involved in target cell killing, in the CAR T cell condition compared to the untransduced T cell control condition alongside increases in IL-6 and sPD-L1/B7-H1 (Fig. 3F). These results suggest specific and robust CAR T cell activity in the on-chip model.

### IL-2 supplementation causes increased target cell killing and release of inflammatory mediators

The cytokine IL-2 is often administered in conjunction with various T cell-based anti-cancer therapies to increase proliferation and persistence of administered cells (Wrangle et al., 2018). To investigate the effect of IL-2 supplementation, co-culture experiments were performed and tumor cell growth in the absence and presence of IL-2 was compared. Addition of 50 IU/mL IL-2 to the endothelial vessel compartment did not affect A375-GFP tumor growth in case of both EpCAM-CD28 CAR T cells and untransduced T cells (Fig. 4A). In contrast, supplementation with IL-2 caused an even stronger response of the EpCAM-CD28 CAR T cells against the HT-29-GFP tumor cells (Fig. 4B). Interestingly, untransduced T cells also showed an anti-tumor response against HT-29-GFP cells in the presence of IL-2 although the effect observed was not as strong as seen in the case of EpCAM-CD28 CAR T cells. To assess the effect of IL-2 supplementation on release kinetics of effector molecules, levels of Granzyme B, IFN-γ, IL-2, IL-6, soluble PD-L1/B7-H1 and TNFα were measured in the supernatant of HT-29-GFP co-cultures after each imaging timepoint (Fig. 4C and D). Without IL-2 supplementation, release of Granzyme B, IFN-γ and IL-2 was constant during the 72-hour co-culture period while both IL-6 and soluble PD-L1/B7-H1 increased readily over time, followed by a drop after the 48-hour timepoint in case of IL-6. In addition, TNFα levels dropped within the first 24 hours and seemed to reach a plateau. In contrast, with IL-2 supplementation, IFN-γ, IL-6, PD-L1/B7-H1 and TNFα readily increased in concentration over time while Granzyme B increased in concentration until 48 hours of co-culture followed by a drop. Interestingly, levels of IL-2 dropped significantly over the course of the co-culture experiment, suggesting either lack of production or consumption of the cytokine. Altogether, these results demonstrate specific and robust CAR T cell activity and showcase the ability of the platform to assess the effects of immunomodulatory compounds like IL-2.

**Figure 4.**
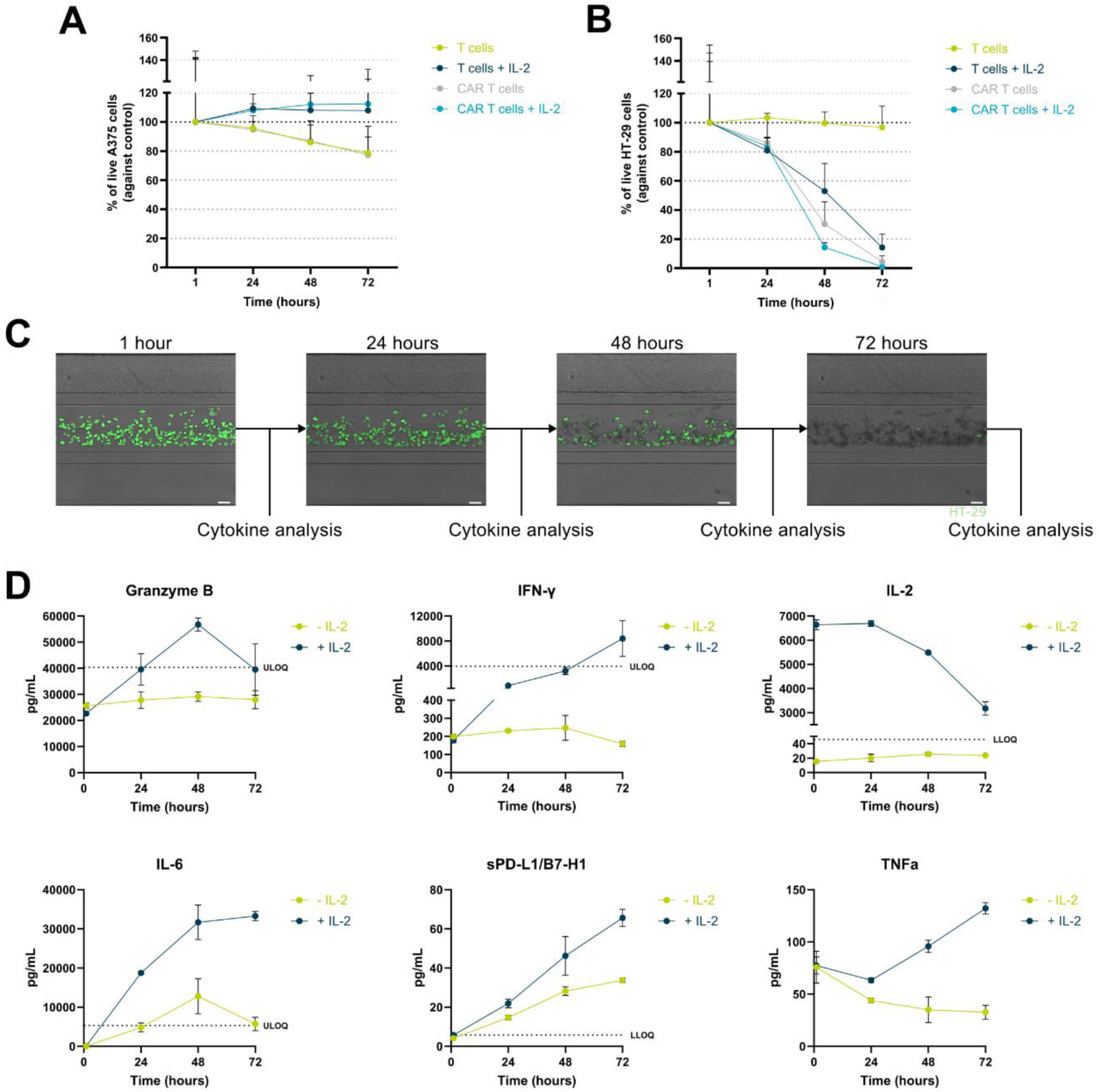
IL-2 supplementation increases HT-29-GFP tumor cell killing and release of inflammatory mediators. (A) Effect of IL-2 supplementation on killing of A375-GFP tumor cells by untransduced T cells and CAR T cells. Data is normalized against the corresponding control condition (+/- IL-2) (n=4). (B) Effect of IL-2 supplementation on killing of HT-29-GFP tumor cells by untransduced T cells and CAR T cells. Data is normalized against the corresponding control condition (+/- IL-2) (n=4). (C) Experimental workflow to assess cytokine release kinetics in co-cultures of HT-29-GFP tumor cells and EpCAM-CD28 CAR T cells. Media samples were gathered after each imaging timepoint and subjected to cytokine analysis using Luminex. (D) Analysis of cytokine levels (pg/mL) in media samples at various timepoint during a 72-hour co-culture of HT-29-GFP tumor cells with EpCAM-CD28 CAR T cells (n=2). Data points outside of lower (LLOQ) and upper (ULOQ) limits of detection (dashed lines) are extrapolated from the standard curve. Data shown in the graphs are presented as mean ± SD. Scale bars indicate 100 µm.

### Killing efficacy of EpCAM-CD28 CAR T cells depends on dosing

One of the risk factors for developing cytokine release syndrome (CRS) is a high infusional dose of CAR T cells (Frey & Porter, 2019). To mimic different dosages, the number of administered CAR T cells was varied resulting in different ratios between the CAR T cells and the HT-29 tumor cells in the culture, called the effector:target (E:T) ratio. As reference, the data that was discussed in the sections above was acquired using 20.000 CAR T cells and 4.000 tumor cells, corresponding to an E:T ratio of 5:1. Killing of HT-29-GFP tumor cells was highly dependent on the number of EpCAM-CD28 CAR T cells added, showing a clear reduction in GFP signal as well as an increasing number of Draq7-positive cells with increasing E:T ratios (Fig. 5A and B). Interestingly, although the lower ratios 1:2, 1:1 and 2:1 were found to affect HT-29-GFP growth when compared to the control condition, the tumor cells did show proliferation after 72 hours of co-culture (223.16±29.21, 174.13±50.21 and 103.44±60.64 respectively) (Fig. 5B). Furthermore, the variability in the anti-tumor response was evident from the range of the error bars. In contrast, HT-29-GFP tumor area decreased during co-culture with E:T ratios of 5:1 and 10:1, where the latter resulted in a stronger anti-tumor effect (21.34±23.26 and 1.13±1.96 respectively after 72 hours). In concordance with the GFP quantification data, the release of effector molecules and inflammatory cytokines in the culture medium was dependent on the E:T ratio as well, showing increasing levels for all tested analytes with increasing numbers of administered EpCAM-CD28 CAR T cells (Fig. 5C). Besides a pronounced dose-dependent effect on the growth of the HT-29-GFP cells and release of effector molecules and inflammatory mediators, the state of the endothelial vessel was also affected by the E:T ratio (Fig. 5A). Looking at vessel morphology after 72 hours, an intact vessel could be seen with E:T ratios of 1:1, 2:1 and 5:1. However, with an E:T ratio of 10:1, the vessel showed clear signs of deterioration. Draq7 indicated increased numbers of dead cells in the endothelial vessel compartment with an increasing E:T ratio (Sup. Fig. 3). Staining for junction marker VE-cadherin after the 72-hour co-culture (Fig. 5D) and subsequent morphometric analysis of the endothelial vessel revealed significant differences between the E:T ratios over a wide range of parameters (Fig. 5E and Sup. Fig. 4). Most notably, confluency was increased for E:T ratios of 1:1, 1:2 and 2:1 and decreased for 5:1 and 10:1 while a change in the Mean Area and Major Axis Angle suggests shrinking and different alignment of the endothelial cells. Altogether, this data highlights the complex relation between the infusion dose and anti-tumor response as well as toxicity. Finally, supplementation with 50 IU/mL IL-2 increased potency of the EpCAM-CD28 CAR T cells for every E:T ratio tested, indicated by a stronger anti-tumor response throughout the co-culture period (Fig. 5F and Sup. Fig. 5A, B and C). The enhanced anti-tumor response to IL-2 supplementation was accompanied by an increase in the release of effector molecules and inflammatory mediators (Fig. 5G and Sup. Fig. 5D).

**Figure 5.**
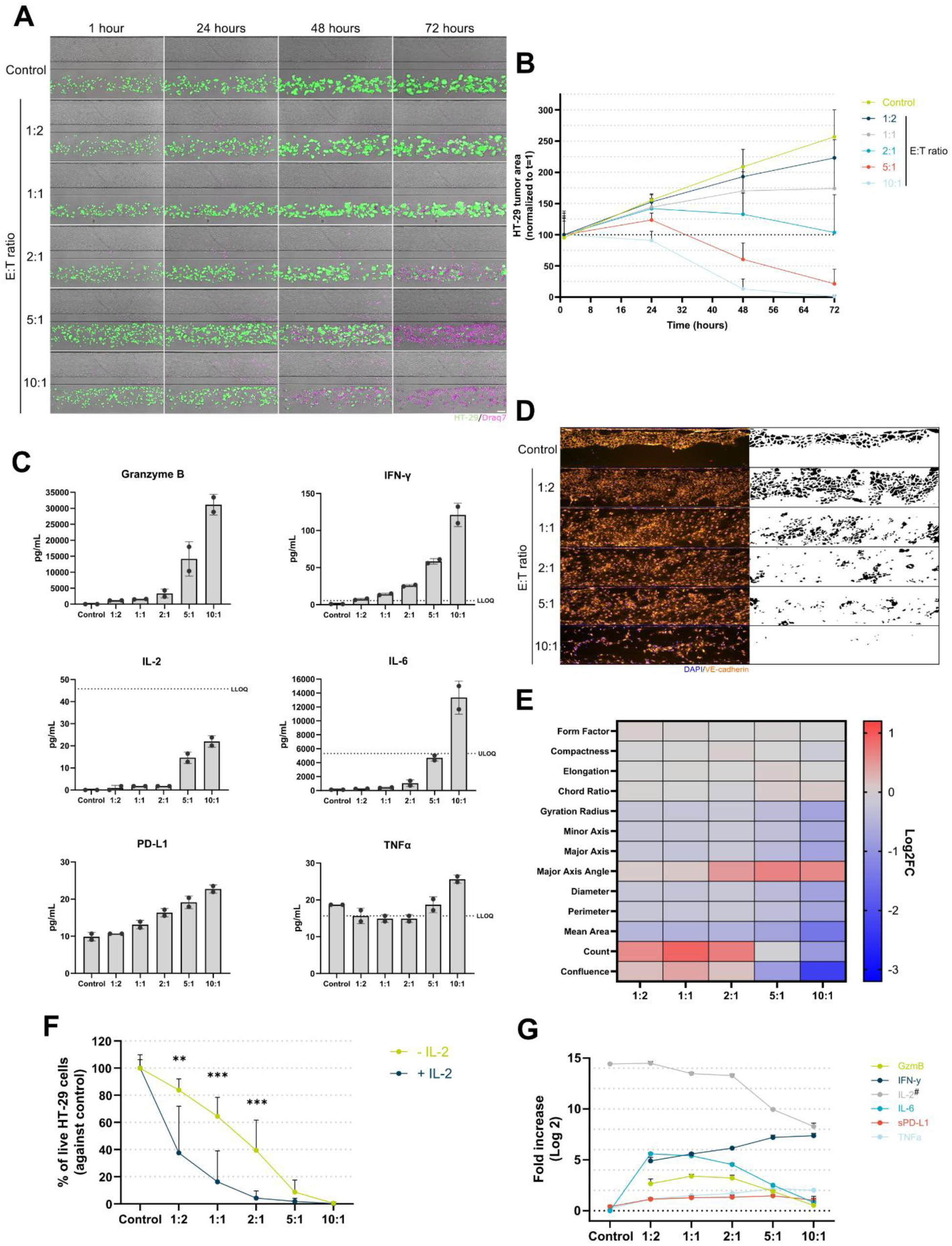
Effect of different E:T ratios on killing of HT-29-GFP tumor cells and endothelial vessel integrity (A) Overview of composite images showing HT-29-GFP tumor cells co-cultured with different numbers of EpCAM-CD28 CAR T cells. Number of CAR T cells used: 1:2 = 2000, 1:1 = 4000, 2:1 = 8000, 5:1 = 20.000 and 10:1 = 40.000. Draq7 signal indicates dead cells. Bottom channels of microfluidic chips are cropped out for visualization purposes. (B) Quantification of HT-29-GFP tumor cell growth during co-culture with EpCAM-CD28 CAR T cells at various E:T ratios. Fluorescent area at each timepoint was normalized against the area at t=1 for each individual replicate (N=2, n=8-10). (C) Analysis of cytokine levels (pg/mL) in media samples after 72 hours of co-culture of HT-29-GFP tumor cells with EpCAM-CD28 CAR T cells at various E:T ratios (n=2). Data points outside of lower (LLOQ) and upper (ULOQ) limits of detection (dashed lines) are extrapolated from the standard curve. (D) Composite maximum intensity images (left) of endothelial vessels that are stained with junction marker VE-cadherin (orange) and nuclei (blue) after 72 hours of co-culture with EpCAM-CD28 CAR T cells at various E:T ratios. Immunofluorescence images were used to create binary masks of intact endothelial cells for morphometric analysis (right). (E) Heatmap showing morphometric parameters measured in the endothelial vessels of HT-29-GFP cultures exposed to EpCAM-CD28 CART cells at different E:T ratios. Data is presented as Log2-transformed fold changes against controls without CAR T cells. (F) Percentage of live HT-29-GFP tumor cells, relative to no CAR T cell controls, at different E:T ratios of EpCAM-CD28 CAR T cells with and without IL-2 supplementation (n=8-13). Statistical analysis was performed using multiple Mann-Whitney tests and FDR correction (**p≤0.01, ***p≤0.001). (G) Effect of IL-2 supplementation on cytokine levels (pg/mL) in media samples after 72 hours of co-culture of HT-29-GFP tumor cells and EpCAM-CD28 CAR T cells at different E:T ratios (n=2). GzmB and IFN-γ fold changes could not be calculated for the control condition. ^#^IL-2 curve was generated using extrapolated values. Data shown in the graphs are presented as mean ± SD.

### Evaluation of EpCAM CAR T constructs with different co-stimulatory domains

The presence of a co-stimulatory domain in the CAR construct ensures activation, proliferation and induction of effector functions of the CAR T cells (Honikel & Olejniczak, 2022). Currently, most CAR T cell products include either a CD28 or 4-1-BB co-stimulatory domain. EpCAM-targeting CAR T cells containing a 4-1-BB co-stimulatory domain, henceforth called EpCAM-4-1-BB CAR T cells, were acquired from the same commercial vendor (Fig. 6A) and the same dosing experiments as described before were performed (Sup. Fig. 6). To compare the killing efficacy of the two CAR T cell products, tumor area was normalized against the no CAR T cell control at each timepoint. While both CAR T cell products resulted in a decrease in HT-29-GFP tumor area with increasing E:T ratios, the EpCAM-CD28 CAR T cells were more potent compared to the EpCAM-4-1-BB CAR T cells (Fig. 6B). Furthermore, not only did the EpCAM-CD28 CAR T cells result in an overall lower percentage of target cells with identical E:T ratios, but these CAR T cells responded to IL-2 supplementation (Fig. 5F) whereas no effect of IL-2 supplementation was observed in case of the EpCAM-4-1-BB CAR T cells (Fig. 6C).

**Figure 6.**
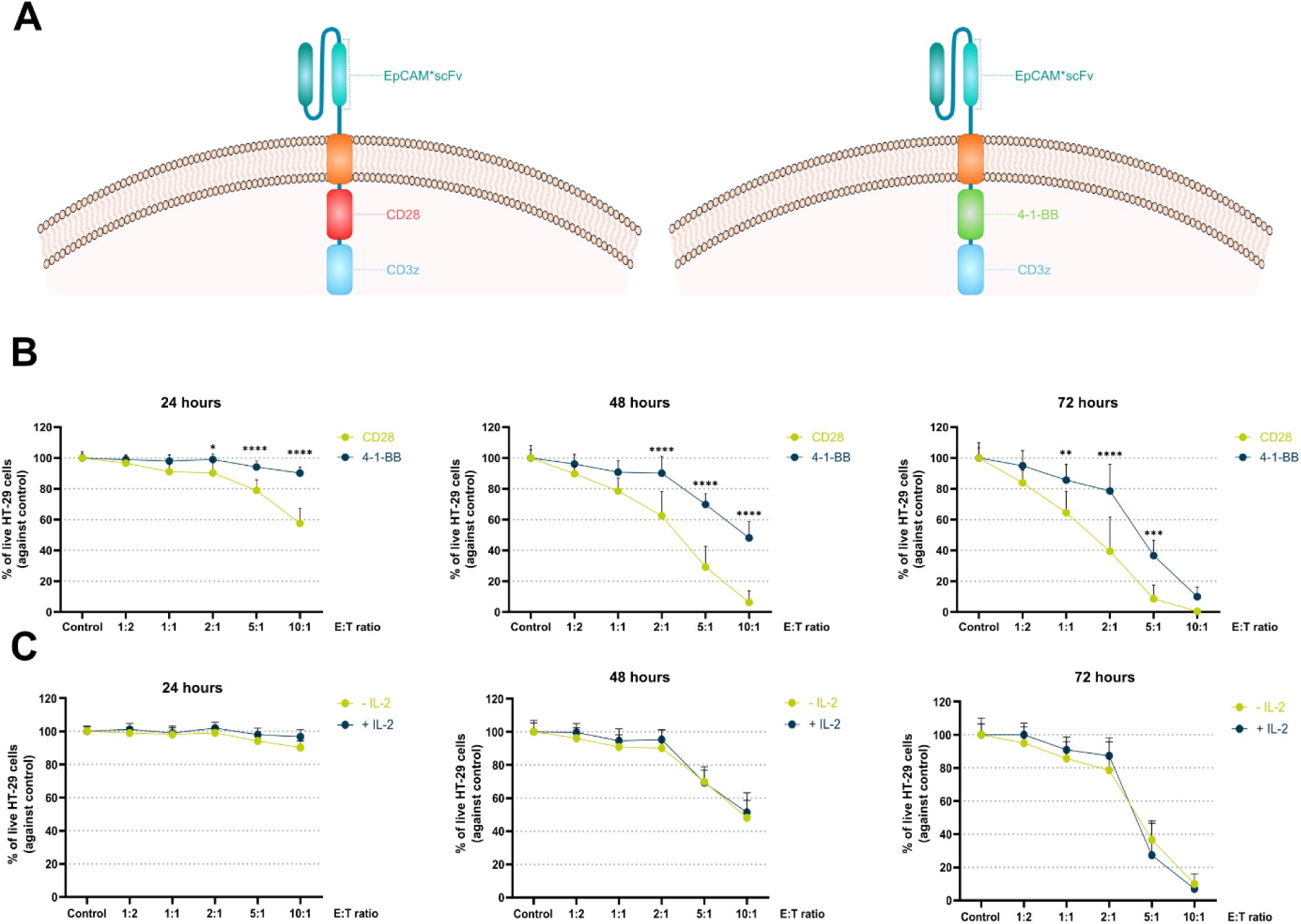
Comparison of EpCAM-CD28 and EpCAM-4-1-BB CAR T cells in killing abilities. (A) Schematic representation of EpCAM-CD28 and EpCAM-4-1-BB CARs. The CAR construct consists of an antigen-binding domain (EpCAM*scFv) fused to a transmembrane domain (orange) and two intracellular signaling domains. Signaling domains of EpCAM-CD28 CAR consists of CD28 (red) and CD3z (blue) while EpCAM-4-1-BB CAR is composed of 4-1-BB (red) and CD3z (blue). (B) Percentage of live HT-29-GFP tumor cells, relative to no CAR T cell controls, after 24 (left), 48 (middle) and 72 (right) hours of co-culture with either EpCAM-CD28 (green line) or EpCAM-4-1-BB CAR T cells (blue line) at different E:T ratios (n=6-10). Statistical analysis was performed using Two-way ANOVA and multiple comparisons (*p≤0.05, **p≤0.01, ***p≤0.001 and ****p≤0.0001). (C) Effect of IL-2 supplementation on killing of HT-29-GFP tumor cells by EpCAM-4-1-BB CAR T cells after 24 (left), 48 (middle) and 72 (right) hours (n=6-7). Data is normalized against the corresponding control condition (+/- IL-2). Data shown in the graphs are presented as mean ± SD.

### Evaluation of clinically relevant treatments in combination with EpCAM-CD28 CAR T cells

Immune checkpoints can be hijacked by tumor cells, preventing clearance of the tumor by sending inhibitory signals to T cells. The use of immune checkpoint inhibitors (ICIs) can reactivate the anti-tumor response by preventing inhibition of the T cells (Fig. 7A). As described earlier, the use of EpCAM-CD28 CAR T at an E:T ratio of 1:1 resulted in a variable anti-tumor response. Therefore, it was investigated whether the response could be enhanced by treatment with ICIs. Since microsatellite stability status affects response to ICIs, experiments were performed with HT-29-GFP (MSS) as well as HCT116-GFP (MSI) tumor cells (Duldulao et al., 2012). HCT116-GFP tumor cells were successfully embedded in the on-chip model and showed a 2-fold proliferation over 72 hours, which is slightly lower compared to HT-29-GFP tumor cells (Sup. Fig. 7A). In addition to Nivolumab (anti-PD-1) and Ipilimumab (anti-CTLA-4), co-cultures were also treated with Temozolomide, an alkylating agent that is reported to sensitize CRC cells to ICIs (25,26). The effect of treatments on the growth of the tumor cells (Sup. Fig. 7B and C) as well as release of Granzyme B, IL-6, soluble PD-L1, and TNFα (Sup. Fig. 8) was assessed. Both the cytokines IL-2 and IFN-γ were found to be under the lower detection limit and therefore omitted from the dataset.

**Figure 7.**
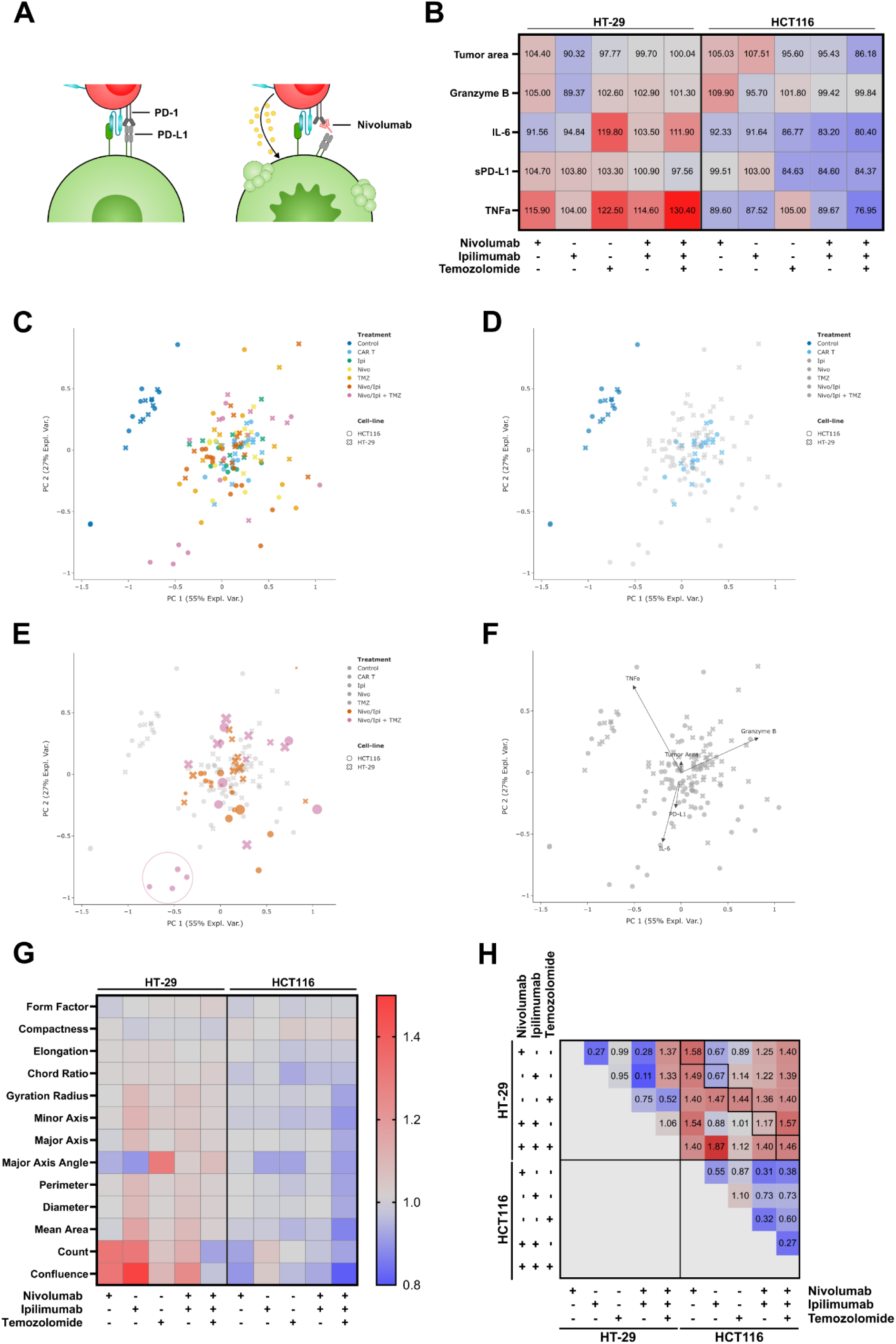
Multiparametric analysis of colorectal cancer cell lines to clinically relevant treatments in combination with EpCAM-CD28 CAR T cells. (A) Schematic representation of the concept of immune checkpoint inhibition. Interaction between PD-1 and PD-L1 is blocked by Nivolumab which restores release of cytotoxic molecules by the CAR T cell (red) (B) Heatmap showing an overview of changes in tumor area and release of Granzyme B, IL-6, sPD-L1 and TNFα in HT-29-GFP and HCT116-GFP co-cultures in response to both single and combination treatments with Nivolumab, Ipilimumab and Temozolomide. Data used to generate the heatmap can be found in supplementary figures 7 and 8. (C,D) PCA plots showing response of HT-29-GFP and HCT116-GFP co-cultures to EpCAM-CD28 CAR T cells and treatments. (E) PCA plot showing the response of HT-29-GFP and HCT116-GFP co-cultures to combination treatments. Datapoints are scaled for the initial tumor cell density at the start of the co-culture. Pink circle indicates a small cluster of replicates consisting of HCT116-GFP cultures exposed to Nivolumab, Ipilimumab and Temozolomide. (F) Contribution and directionality of the different features of the principal components. (G) Heatmap showing morphometric parameters measured in the endothelial vessels of HT-29-GFP and HCT116-GFP co-cultures exposed to EpCAM-CD28 CART cells and treatments. Shown are fold changes against untreated controls. (H) Correlation distance matrix showing similarity between treatment responses based on patterns in relative change in tumor area, cytokine release and endothelial response. Data are presented as unsquared correlation distances, with values <1 indicating similarity in response while values >1 indicate distinct responses. A value of 1 means no relation between observed responses and a value of 0 indicates an identical response.

Although the treatments did not decrease variability in response to the CAR T cells at a low E:T ratio, some changes in average tumor area and cytokine levels were seen (Fig. 7B). Most notably, average tumor area of HCT116-GFP cells decreased in response to treatment with a combination of Nivolumab, Ipilimumab and Temozolomide (13.82% decrease) while HT-29-GFP cells were unaffected. Furthermore, levels of IL-6, sPD-L1 and TNFα were also affected differently in response to this combination treatment. Principal component analysis (PCA) was performed to gain more insight into the variable responses to the treatments (Fig. 7C). HCT116 (blue dots) and HT-29 (blue crosses) control chips without CAR T cells formed a distinct cluster, clearly separating from co-cultures with CAR T cells (Fig. 7D, light blue). No clear treatment-specific or tumor cell line-specific clustering was observed in case of single treatment conditions with either Nivolumab, Ipilimumab or Temozolomide (Sup. Fig. 9E). However, in case of combination treatments, a small cluster consisting of four datapoints could be seen that exclusively contained replicates of HCT116 co-cultures exposed to combination treatment with Nivolumab, Ipilimumab and Temozolomide (Sup. Fig. 9G). Scaling the data points for the initial density of the tumor cells revealed that the replicates in this small cluster had a lower number of tumor cells at the start of the co-culture with CAR T cells compared to the other replicates from the same treatment condition (Fig. 7E, pink circled cluster). No clear relation between tumor cell density and clustering was found for other treatment conditions or HT-29-GFP co-cultures (Sup. Fig. 9B, F and H). Contribution and directionality of the features to the principal components was assessed (Fig. 7F). As expected, Granzyme B drives separation of the tumor cell only controls from the co-cultures with CAR T cells. Furthermore, the small cluster that was described before was characterized by a decrease in tumor area after 72 hours (38.41% compared to control), indicating an effect of the treatment on the growth of the HCT116 tumor cells (Sup. Fig. 10). In addition, levels of IL-6 (60.11%), sPD-L1 (37.57%) and TNFα (79.68%) were also strongly decreased. Altogether, these results suggest an effect of combination treatment with Nivolumab, Ipilimumab and Temozolomide that is specific for HCT116-GFP cells and is dependent on the number of tumor cells present at the start of co-culture with CAR T cells.

Morphometric analysis of the VE-cadherin network was conducted to assess stability of the endothelial vessel and potential signs of toxicity in response to treatment. Endothelial vessels were found to be similarly affected by addition of CAR T cells in untreated co-cultures with either HT-29-GFP or HCT116-GFP tumor cells (Sup. Fig. 11). However, clear differences could be seen across a wide range of descriptors when co-cultures were treated (Fig. 7G). Not only did the different treatments cause distinct changes to the endothelial vessel within the HT-29 and HCT116 groups, but the same treatment also seemed to affect the endothelial vessel differently depending on co-culture with HT-29 or HCT116.

Unsquared correlation distance was used to assess similarity between treatment responses based on patterns in the relative change in tumor area, cytokine release and endothelial response (Fig. 7H). Similar responses are indicated by correlation distances <1 while distinct treatment effects are indicated by correlation distances >1, with progressively lower and higher values indicating respectively higher similarity and dissimilarity. Lastly, a correlation distance of 1 indicates no relation between treatment conditions. Most notably, combined treatment with Nivolumab, Ipilimumab and Temozolomide caused a distinct response of HT-29 co-cultures compared to other treatments, indicated by correlation distances ranging from 1.06 to 1.37, with exception of treatment with Temozolomide (0.52). In contrast, these treatments all caused similar responses in HCT116 co-cultures (0.27 to 0.73). Pairwise comparison between responses of HT-29-GFP and HCT116-GFP co-cultures, indicated by the boxed values, confirmed that response to treatment differs between HT-29-GFP and HCT116-GFP cultures in case of Nivolumab (1.58), Temozolomide (1.44), Nivolumab + Ipilimumab (1.17) as well as Nivolumab + Ipilimumab + Temozolomide (1.46). However, response was somewhat similar in case of treatment with Ipilimumab (0.67). Collectively, the data presented in this figure demonstrates how the model can be utilized to investigate clinically relevant compounds and mechanisms using a multiparametric approach.

## 4. Discussion

In this study, we described an on-chip model comprising endothelium and ECM-embedded tumor cells to evaluate different aspects of CAR T cell activity, e.g. infiltration, recognition and killing of target cells and secretion of effector molecules. We demonstrated specific and robust killing of target cells using a semi-automated quantification method based on GFP expression, showing a clear reduction in EpCAM-positive HT-29-GFP tumor cells in presence of CAR T cells and a concomitant release of effector molecules such as Granzyme B and IFN-γ. Moreover, both the anti-tumor response and cytokine release could be enhanced by IL-2 supplementation.

A major challenge in CAR T cell therapy for solid tumors is the lack of sufficient infiltration, causing CAR T cells to be challenged with a great number of tumor cells per CAR T cell which demands extremely high activity and persistence (10). Therefore, the infusion dose is usually maximized to improve infiltration. However, this increases risk of developing CRS, meaning the patient suffers from systemic inflammation due to high levels of pro-inflammatory cytokines that are released (27). Several experiments were performed using varying numbers of CAR T cells, resulting in distinct E:T ratios and resembling different doses. We observed a dose-dependent effect on the growth of the target cells and corresponding release of effector molecules and other pro-inflammatory cytokines. Furthermore, whereas integrity of the endothelial vessel improved with addition of CAR T cells at low E:T ratios, higher E:T ratios caused disruption of the vessel, likely due to increased levels of pro-inflammatory cytokines that were released by the CAR T cells and were measured in our study. The model therefore successfully captures interactions of the CAR T cells with both target and bystander cells, allowing the possibility to measure and study potential mediators of CAR T cell-associated adverse events involving bystander cells, like endothelial toxicity (28).

We then utilized the model to evaluate two commercially available CAR T constructs that contain different co-stimulatory domains. All six FDA-approved CAR T cell products contain either a CD28 (Yescarta & Tecartus) or 4-1-BB (Kymriah, Carvykti, Breyanzi & Abecma) co-stimulatory domain. EpCAM-CD28 CAR T cells were more potent compared to EpCAM-4-1BB CAR T cells in our on-chip model. Various studies have been conducted to try and compare CAR T cells with distinct co-stimulatory domains, revealing differences in effector function, survival and safety profiles (29–31). In vivo, CAR T cell products containing the 4-1-BB co-stimulatory domain are generally reported to result in more robust anti-tumor responses against hematological malignancies (31,32). However, these differences in potency are due to higher persistence of the 4-1-BB CAR T cells in the patients’ body after administration, something that is not considered in our model. In fact, CAR T cells containing a CD28 co-stimulatory domain have been found to exhibit faster proliferation and stronger signaling as well as higher rates of toxicity (Salter et al., 2018). We also observed a different response to IL-2 supplementation between the two CAR T cell products. Whereas the killing capacity of EpCAM-CD28 CAR T cells increased, EpCAM-4-1-BB CAR T cells did not seem to respond to IL-2. Although the exact composition is undisclosed, AIM-V medium that is used for the co-cultures is known to contain IL-2. We therefore hypothesize that the IL-2 in the medium already maximizes the killing potential of the EpCAM-4-1-BB CAR T cells, leaving them unresponsive to further increases in IL-2. In addition, co-stimulation of CD28, but not 4-1-BB, is reported to confer resistance to TGFβ repression via IL-2 which could contribute to the observed differences (34).

Since effective CAR T cell therapy for solid tumors proves to be challenging, potential synergy between CAR T cells and ICIs is being investigated (35). We utilized the model to test clinically relevant treatments to see if CAR T cell-mediated tumor cell killing at a low E:T ratio could be enhanced. Perhaps unsurprising considering clinical observations, ICIs showed limited effect on growth of both HT-29-GFP and HCT116-GFP tumor cells. However, based on patterns in change in tumor area, cytokine levels and endothelial integrity, it was found that co-cultures responded differently to treatment depending on the presence of HT-29-GFP or HCT116-GFP cells. Notably, combination treatment with Nivolumab, Ipilimumab and Temozolomide caused reduction in HCT116-GFP tumor area in a subset of replicates that contained relatively low numbers of tumor cells at the start of the co-culture, which likely created a more favorable setting for the CAR T cells. This exemplifies how crucial sufficient infiltration of effector cells is to generate a robust response but also demonstrates the sensitivity of our on-chip model. Interestingly, this response was also characterized by lowered levels of IL-6 and soluble PD-L. Both are described as prognostic indicators of clinical outcome, with lower levels being associated with better prognosis and decreased occurrence of adverse effects after treatment with ICIs (36–38).

Since HCT116-GFP tumor cells are MMR deficient, the observed response to Temozolomide in combination with Nivolumab and Ipilimumab is surprising (39). Furthermore, it is also unlikely that the observed response is due to hypermutation and increased TMB after a single dose of Temozolomide considering its short half-life of 1.8 hours (40). Therefore, it seems that there might be a secondary mechanism at play that causes increased sensitization of HCT116 to ICIs and CAR T cell-mediated killing which requires further investigation. Nonetheless, this dataset demonstrates clearly how the model can be used to investigate clinically relevant compounds and mechanisms using a multiparametric approach by leveraging the throughput and accessibility of the platform.

Most models and assays currently used to study CAR T functionality generally focus on a single aspect, e.g. migration potential in response to chemokines in a Transwell set up or cytokine release in response to target cell recognition (11). However, several MPS models that aim to study CAR T functionality in a more complex *in vivo*-like setting have been developed recently. Despite increased biological relevance, these systems often require specific equipment, making them tedious to establish (16,41). Furthermore, inclusion of vasculature is often not straightforward and is sometimes achieved by seeding vascular endothelial cells as a monolayer on a membrane (15). Tumor vasculature acts as a physical barrier for CAR T cell infiltration and has more than once been suggested as a potential therapeutic target (42). Furthermore, endothelial cells are actively involved in shaping immune response as they can express several inhibitory molecules, among which PD-L1 (43). Finally, immunotherapies can cause cardiovascular adverse events like endothelial dysfunction (28,44). Altogether, this highlights the importance of including endothelial cells in preclinical models. Our model includes a tubular endothelial vessel through which CAR T cells are perfused, adjacent to ECM-embedded tumor cells. Therefore, migration that is displayed by the CAR T cells is not driven by gravity and the lack of an artificial membrane allows for unimpeded extravasation and infiltration. Moreover, most of the abovementioned MPS are not scalable, making dosing experiments and exposure to compound combinations difficult, laborious and costly. The platform used in this study contains 40 individual microfluidic chips per plate, allowing for a higher throughput while maintaining biological complexity. In addition, previous research already highlighted automation compatibility of the OrganoPlate platform, allowing mid-to high-throughput screening experiments (45). Finally, mass manufacturing and established QC methods make the platform ready for clinical implementation. Therefore, to our knowledge, the model presented in this study allows preclinical *in vitro* studies on CAR T functionality at unprecedented detail and scale.

However, the current model does have some limitations. Being limited to commercially available material, the endothelial cells and CAR T cells are not donor-matched meaning the CAR T cells could potentially recognize the endothelial cells in an MHC-dependent manner (46). This possibly contributes to the observed disruption of the endothelial vessel. Moreover, this also made it impossible to include suppressive cells like tumor-associated macrophages (TAMs) in our model, although these affect clinical outcomes (47). In future projects, we hope to utilize patient-derived material to generate a donor-matched culture.

## 5. Conclusion

In summary, we developed an on-chip model comprising an endothelial vessel and ECM-embedded tumor cells and demonstrated infiltration of CAR T cells as well as robust and specific dose-dependent target cell killing. We believe the model presented in this study provides researchers with a powerful and versatile tool to study crucial aspects of CAR T functionality in the context of solid tumors, all in one platform. However, we envision incorporation of suppressive cells like TAMs to improve the model’s biological value even further. Lastly, the model is not limited to CAR T cells but can also be employed to study other cellular therapies like CAR NK and TILs.

## Acknowledgements

We would like to thank the MIMETAS BioCore team for providing us with the OrganoReady^®^ Lumenised Collagen 3-lane 40 product. We would also like to thank Frederik Schavemaker for his help in designing schematics. This research was supported by Oncode Accelerator, a Dutch National Growth Fund project under grant number NGFOP2201.

## Author contributions

Conceptualization: LH, JS, CA, LB and KQ

Methodology: LH, AO, TO, JW, JS and KQ

Data curation and analysis: LH, AO, TO and JW

Supervision: TB, LB and KQ

Writing-draft preparation: LH, TO, JW and KQ

Writing-review and editing: all authors

## Supplementary figures

**Supplementary Figure 1.**
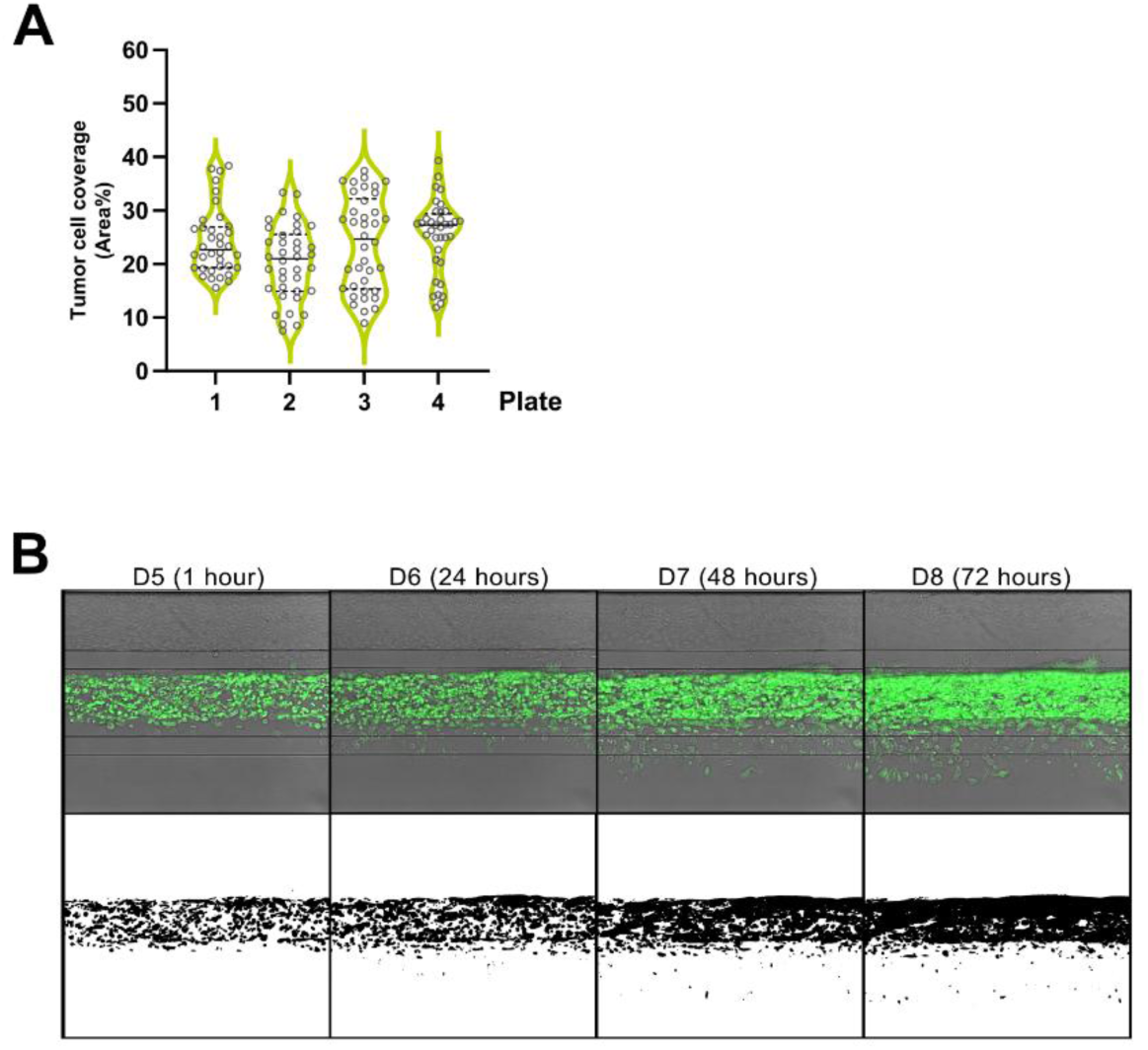
Quantification of HT-29-GFP area across plates and generation of an EpCAM-negative tumor compartment using A375-GFP tumor cells (A) Variability of the HT-29-GFP tumor cell seeding was assessed by calculating the tumor cell coverage of the middle channel, indicating an average Area% of 23.41±7.49 based on 148 microfluidic chips across 4 different OrganoPlates. Data included in the graphs are presented as median and 1^st^ and 3^rd^ quartiles (dashed lines). (B) A375-GFP tumor cells proliferate after introduction in the lumenised ECM and cause regression of the endothelial vessel over time. Like HT-29-GFP cells, the fluorescent signal could be used to create a binary mask that was used to track growth of A375-GFP tumor cells.

**Supplementary Figure 2.**
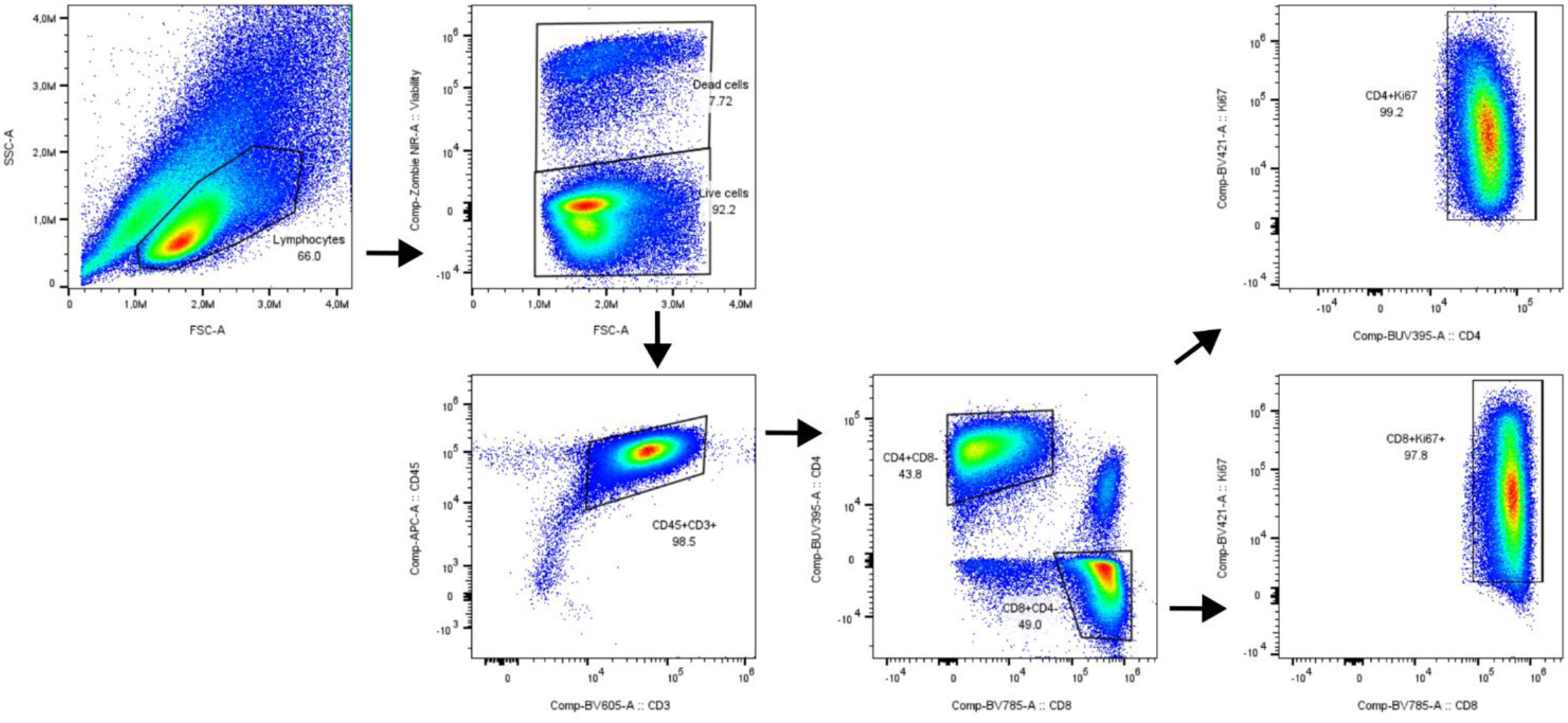
Flow cytometry assessment of Human EpCAM scFv-TM-CD28-CD3ζ CAR T cells. Live CAR T cells (CD45+CD3+) were gated into CD4+CD8- (helper T cells) and CD4-CD8+ (cytotoxic T cells) populations to determine CD4+/CD8+ ratio. The proliferative state of both CD4+CD8- and CD4-CD8+ populations was determined by the functional marker Ki-67. Gating for Ki-67 positive cells was based on FMO control.

**Supplementary Figure 3.**
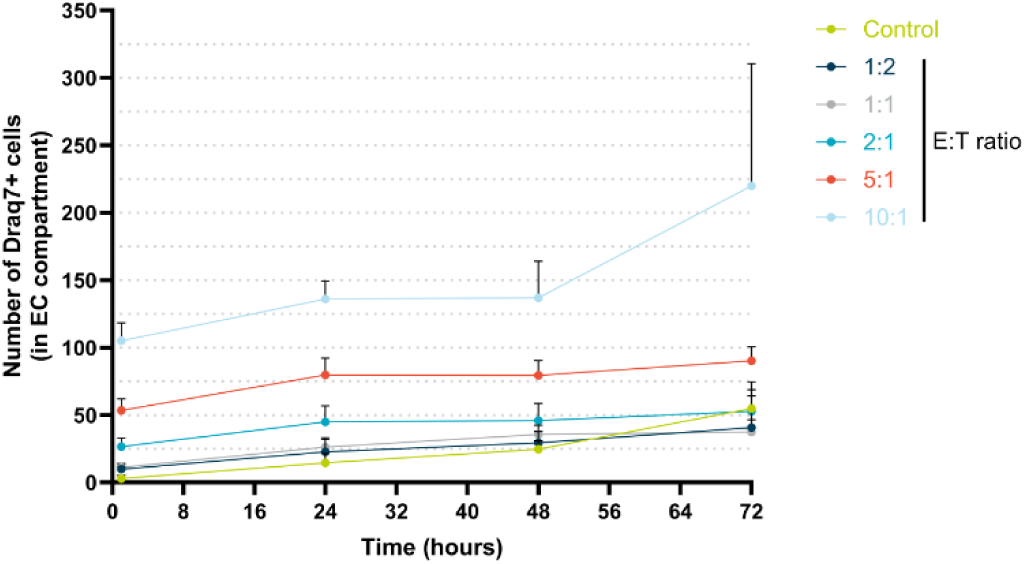
Quantification of dead cells in the endothelial vessel in presence of EpCAM-CD28 CAR T cells at different E:T ratios. Co-culture medium was supplemented with Draq7 to visualize dead cells. The number of Draq7-positive cells was quantified during 72 hours of co-culture with EpCAM-CD28 CAR T cells at various E:T ratios (n=6-8). Data are presented as mean ± SD.

**Supplementary Figure 4.**
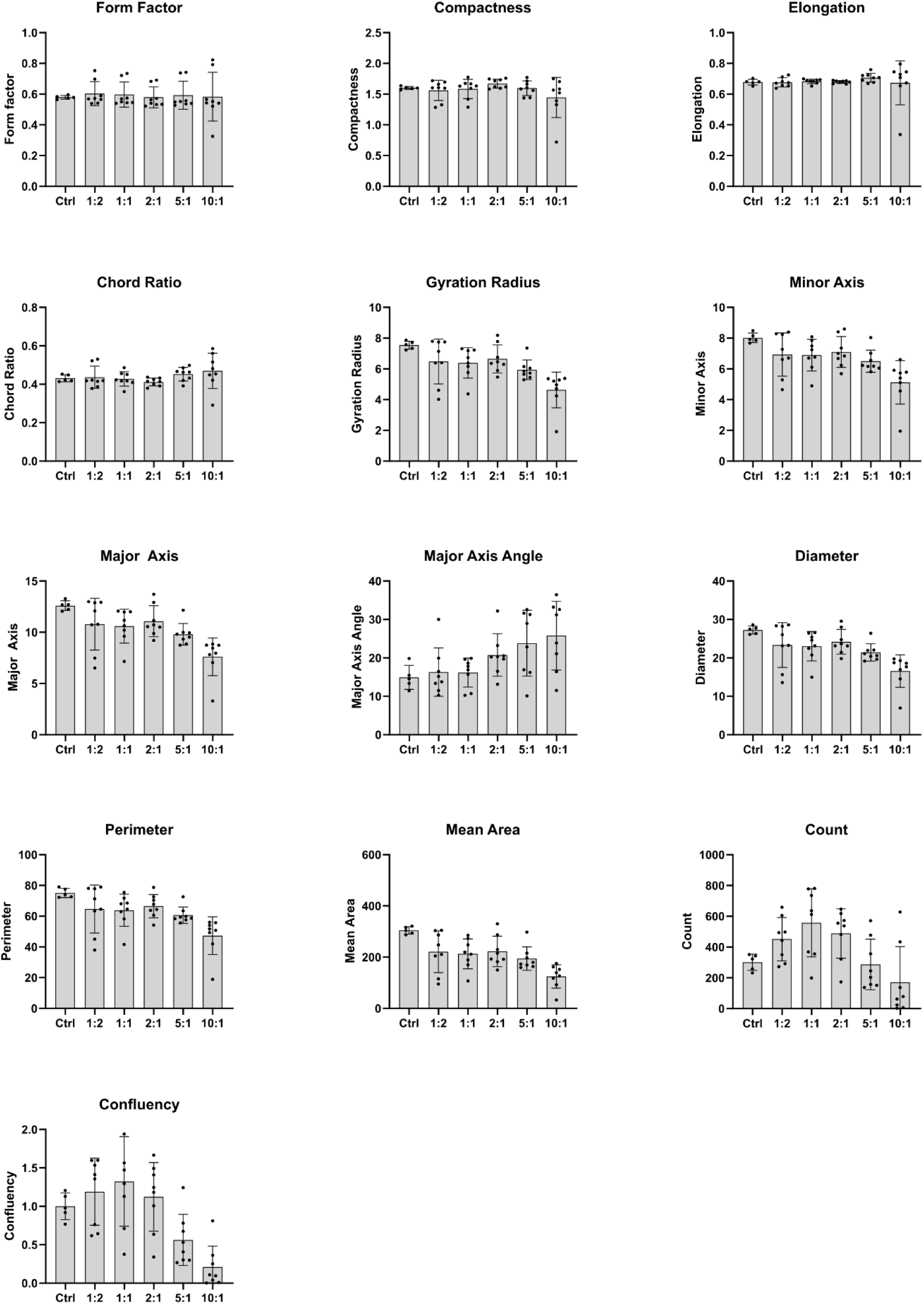
Morphometric analysis of endothelial vessels co-cultured with EpCAM-CD28 CAR T cells at different E:T ratios. Endothelial vessels were exposed to EpCAM-CD28 CAR T cells at different E:T ratios (1:2, 1:1, 2:1, 5:1 and 10:1) and stained with VE-cadherin after the 72-hour co-culture. The endothelial response was assessed by analysis of VE-cadherin objects using a range of different morphological and spatial descriptors using IN Carta® (n=5-8). Data are presented as mean ± SD.

**Supplementary Figure 5.**
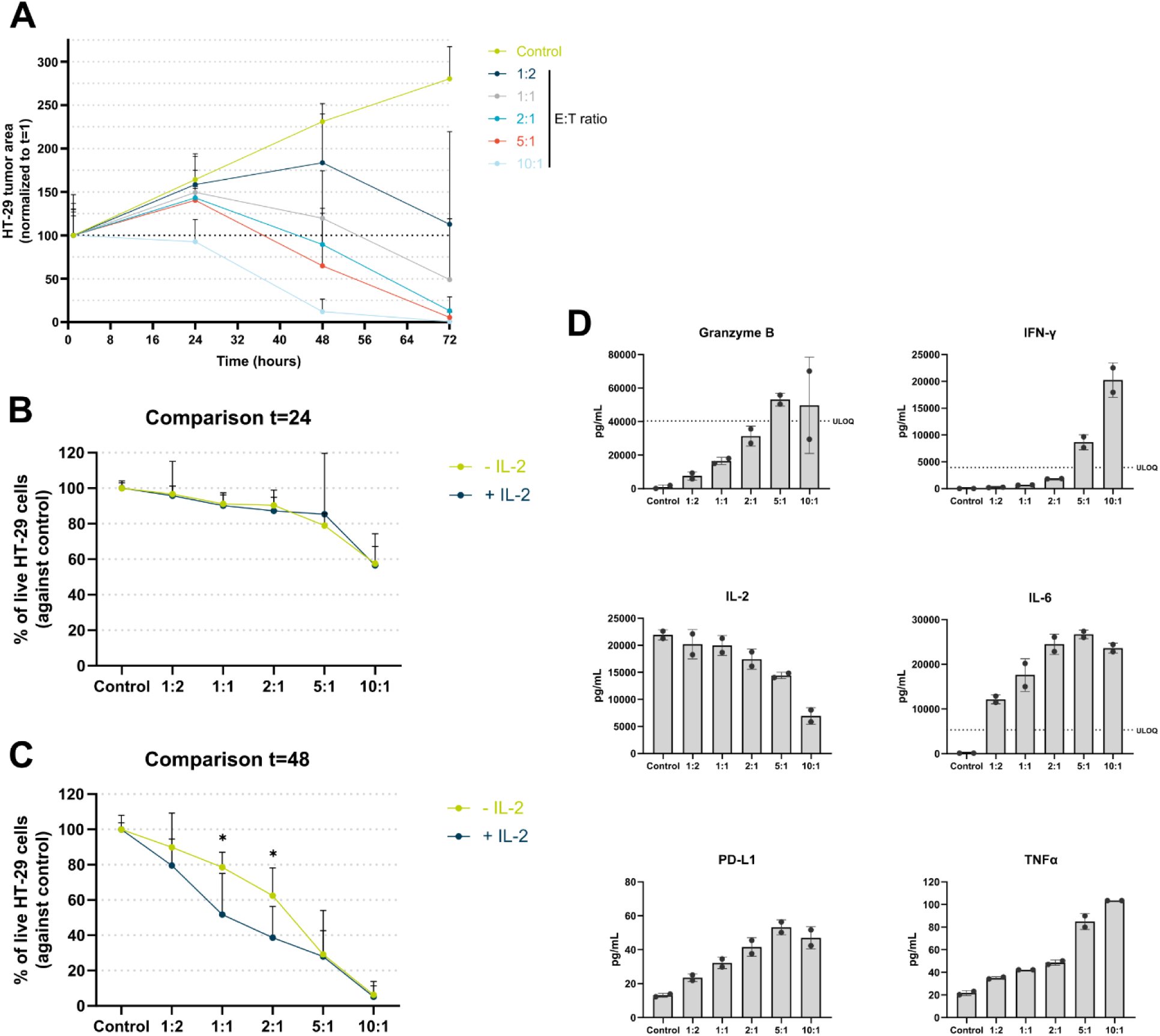
Effect of IL-2 supplementation on CAR T-cell mediated target cell killing and release of effector molecules. (A) Quantification of HT-29-GFP tumor cell growth during co-culture with EpCAM-CD28 CAR T cells at various E:T ratios in the presence of IL-2. Fluorescent area at each timepoint was normalized against the area at t=1 for each individual replicate (N=2, n=10-12). (B,C) Effect of IL-2 supplementation on killing of HT-29-GFP tumor cells by EpCAM-CD28 CAR T cells at different E:T ratios (1:2, 1:1, 2:1, 5:1 and 10:1) after co-culture for 24 (B) and 48 hours (C) (N=2, n=8-12). (D) Analysis of cytokine levels (pg/mL) in media samples after 72 hours of co-culture of HT-29-GFP tumor cells with EpCAM-CD28 CAR T cells at various E:T ratios in the presence of IL-2 (n=2). Values are normalized against respective control conditions with or without IL-2. Data included in the graphs are presented as mean ± SD.

**Supplementary Figure 6.**
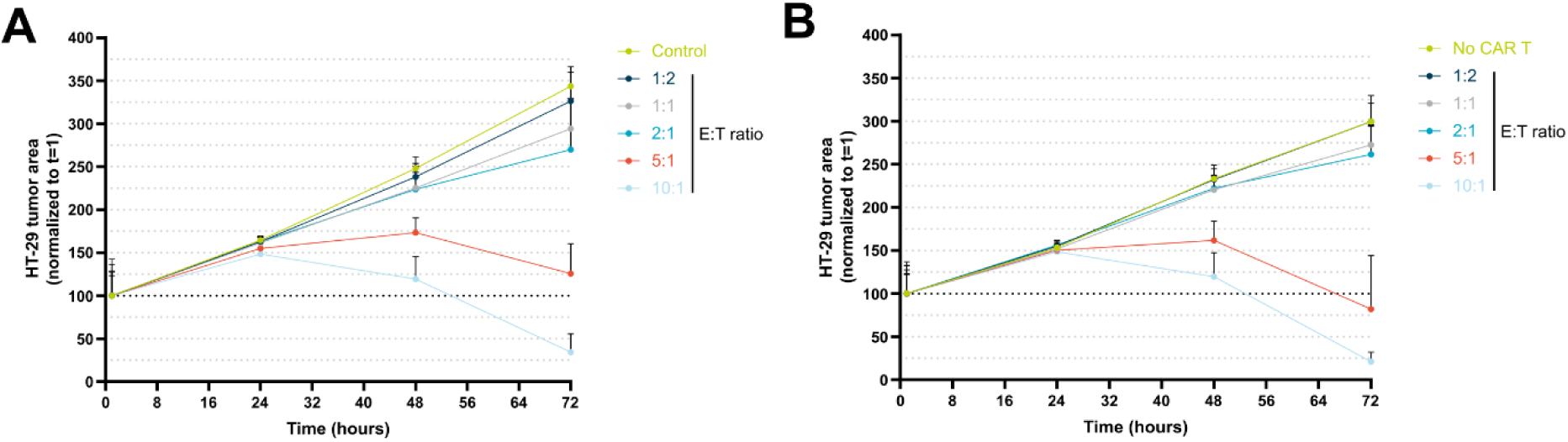
Effector:target experiments with EpCAM-4-1BB CAR T cells. (A) Quantification of HT-29-GFP tumor cell growth during co-culture with EpCAM-4-1-BB CAR T cells at various E:T ratios. Fluorescent area at each timepoint was normalized against the area at t=1 for each individual replicate (n=6-7). (B) Quantification of HT-29-GFP tumor cell growth during co-culture with EpCAM-4-1-BB CAR T cells at various E:T ratios in the presence of IL-2. Fluorescent area at each timepoint was normalized against the area at t=1 for each individual replicate (n=6-7). Data included in the graphs are presented as mean ± SD.

**Supplementary Figure 7.**
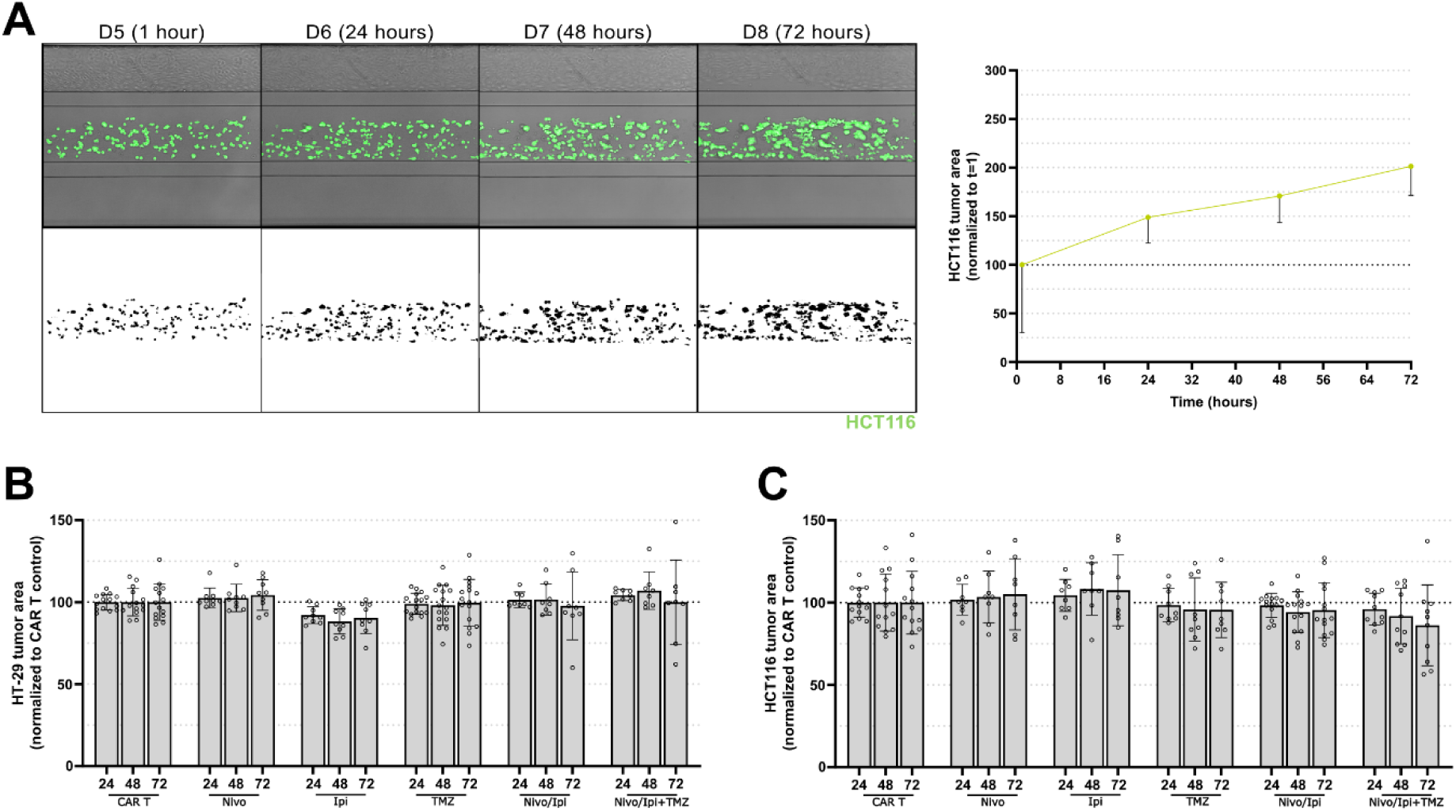
Quantification of HCT116-GFP tumor area and effect of treatment with ICIs and Temozolomide on tumor growth. (A) HCT116-GFP tumor cells proliferate after introduction in the lumenised ECM and cause regression of the endothelial vessel over time. The fluorescent signal could be used to create a binary mask that was used to track growth of HCT116-GFP tumor cells from day 5 until day 8. Fluorescent area at each timepoint was normalized against the area at t=1 for each individual replicate (n=15). (B) Effect of treatments on HT-29-GFP tumor area. Data was normalized against the untreated control at each timepoint (n=8-16). (C) Effect of treatments on HCT116-GFP tumor area. Data was normalized against the untreated control at each timepoint (n=8-14). Data included in the graphs are presented as mean ± SD.

**Supplementary Figure 8.**
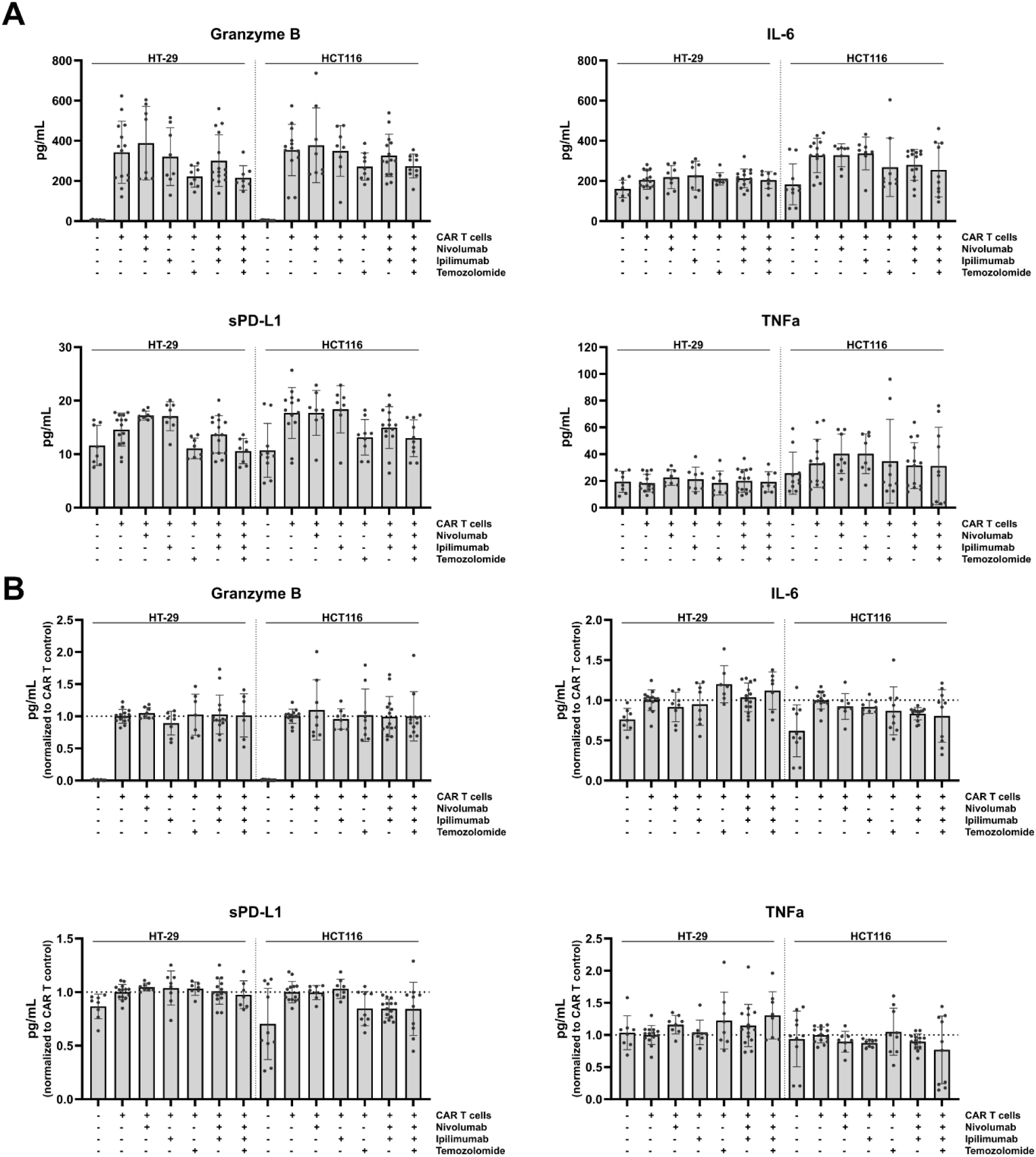
Effect of treatment with ICIs and Temozolomide on cytokine release. (A) Analysis of cytokine levels (pg/mL) in media samples after 72 hours of co-culture of HT-29-GFP and HCT116-GFP tumor cells with EpCAM-CD28 CAR T cells. Co-cultures were untreated or exposed to Nivolumab, Ipilimumab and/or Temozolomide (n=8-14). (B) Cytokine levels in panel A were normalized against untreated controls per experiment resulting in fold changes that were used for principal component analysis (n=8-14). Data included in the graphs are presented as mean ± SD.

**Supplementary Figure 9.**
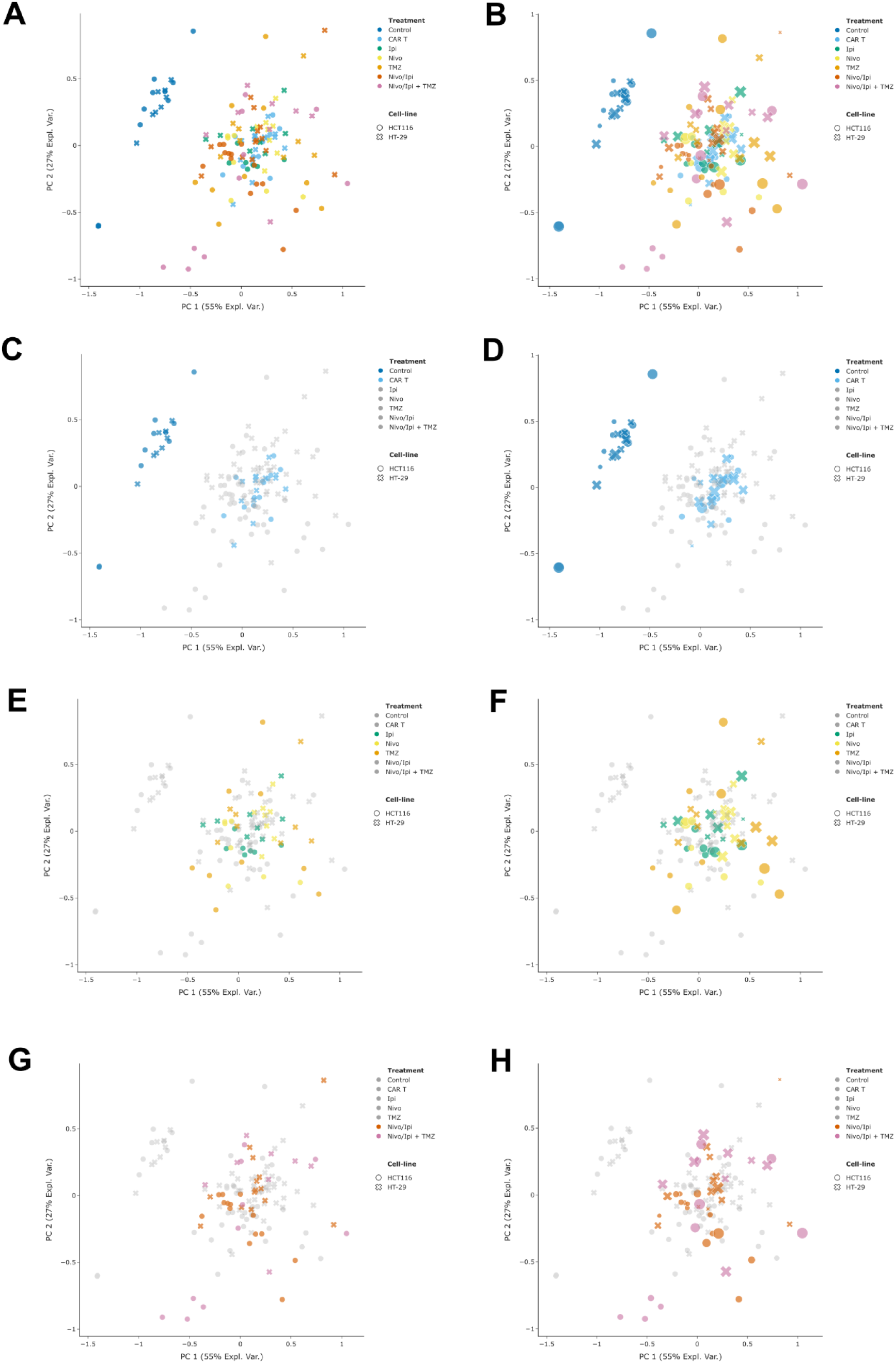
Principal component analysis (PCA) visualizations of treatment responses. Cytokine levels of Granzyme B, IL-6, PD-L1/B7-H1 and TNFα as well as normalized tumor area based on GFP signal after 72 hours of co-culture were combined for these plots. Left panels show PCA visualization with all included features, right panels show the same plots where datapoints are scaled to the initial tumor cell density at the start of the co-culture. HCT116 co-cultures are indicated with circles while HT-29 co-cultures are indicated with crosses. Treatments are color-coded. (A,B) Full visualization of untreated co-cultures and all treatments. (C,D) Highlight of control conditions. Untreated controls with and without EpCAM-CD28 CAR T cells show clear separation among the x-axis (PC1). (E,F) Highlight of single treatment with Nivolumab, Ipilimumab, or Temozolomide. No clear grouping is visible for the two tested cell lines. (G,H) Highlight of combination treatments (Nivolumab & Ipilimumab with or without Temozolomide). Four datapoints with a relatively low seeding density separate from the main cluster.

**Supplementary Figure 10.**
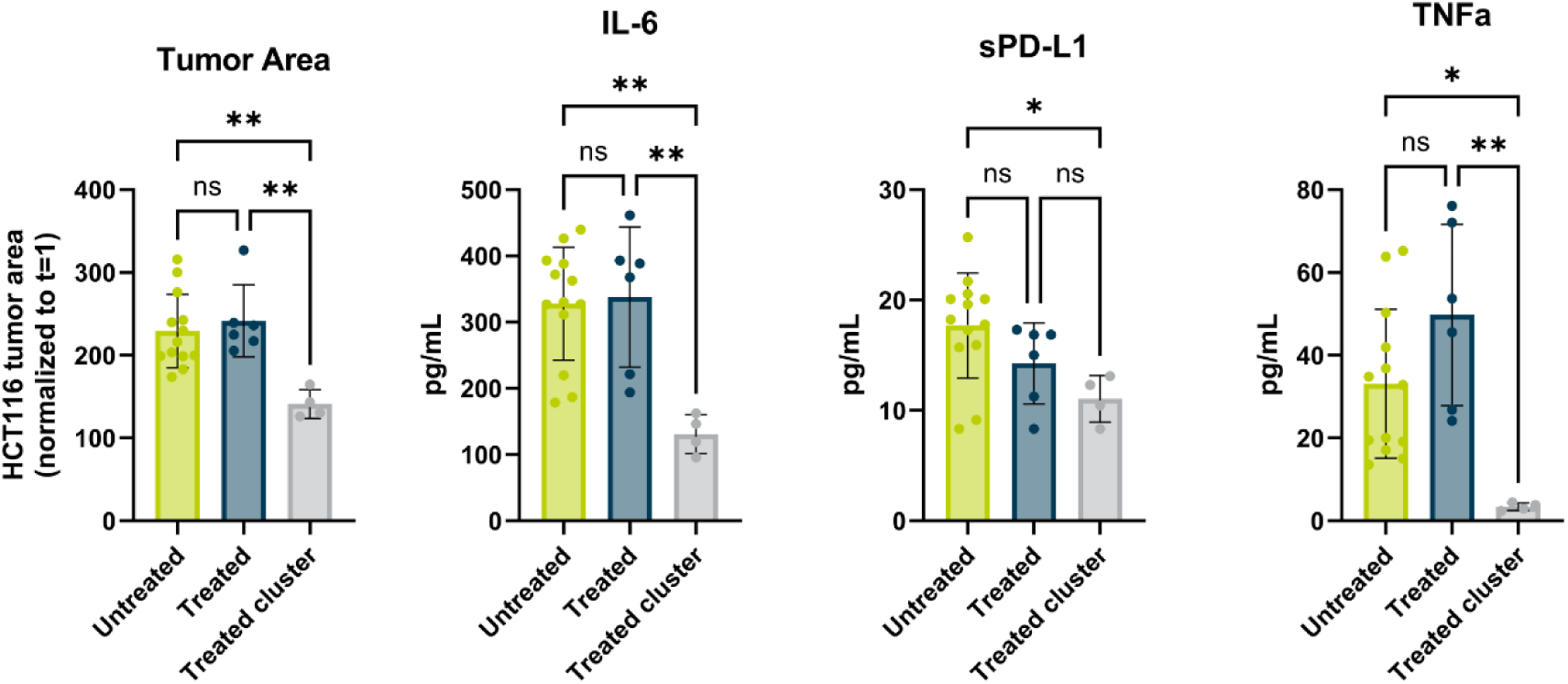
Subset of HCT116-GFP co-cultures responds to treatment with Nivolumab, Ipilimumab and Temozolomide. Exposure to combination treatment with Nivolumab, Ipilimumab and Temozolomide causes decreases in HCT116 tumor area (left) and release of IL-6 (middle left), sPD-L1 (middle right) and TNFa (right) in a subset of replicates (Treated cluster, n=4). Exposed but non-responding group is annotated by ‘Treated’ (n=6). Statistical analysis was performed using One-way ANOVA and multiple comparisons (*p≤0.05, **p≤0.01). Data included in the graphs are presented as mean ± SD.

**Supplementary Figure 11.**
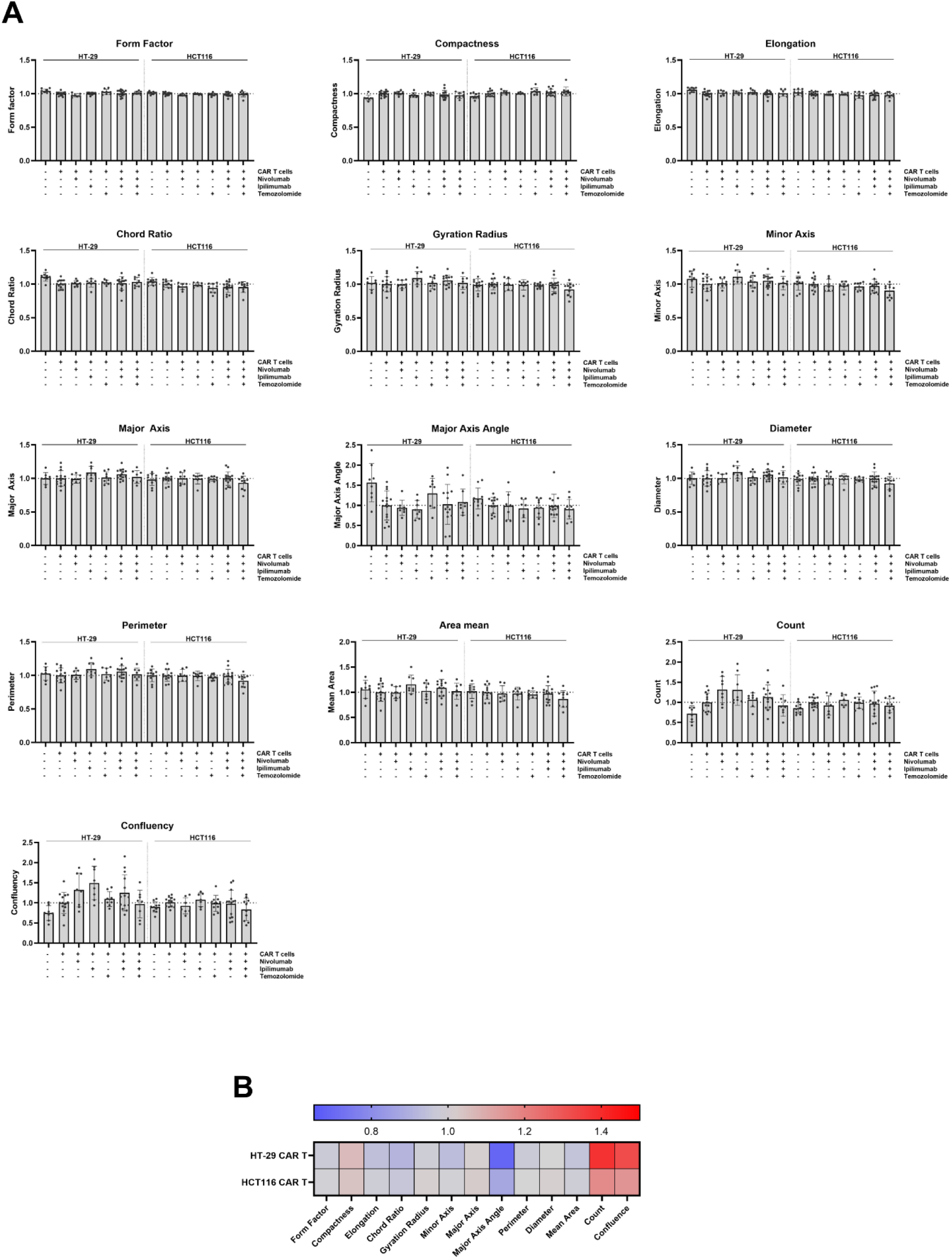
Morphometric analysis of endothelial vessels co-cultured with either HT-29 or HCT116 tumor cells. (A) Endothelial vessels were co-cultured with either HT-29-GFP or HCT116-GFP tumor cells and stained with VE-cadherin after the 72-hour co-culture. Co-cultures were either untreated or exposed to single or combination treatments with Nivolumab, Ipilimumab and Temozolomide in the presence of EpCAM-CD28 CAR T cells (E:T = 1:1). The endothelial response was assessed by analysis of VE-cadherin objects using a range of different morphological and spatial descriptors using IN Carta® (n=7-14) and normalization against respective untreated controls containing CAR T cells for each parameter. Data are presented as mean ± SD. (B) Heatmap showing change in morphometric parameters measured in endothelial vessels, co-cultured with either HT-29-GFP or HCT116-GFP tumor cells, upon addition of EpCAM-CD28 CAR T cells (E:T = 1:1). Data is presented as fold changes against respective controls without CAR T cells.

**Supplementary table 1.**
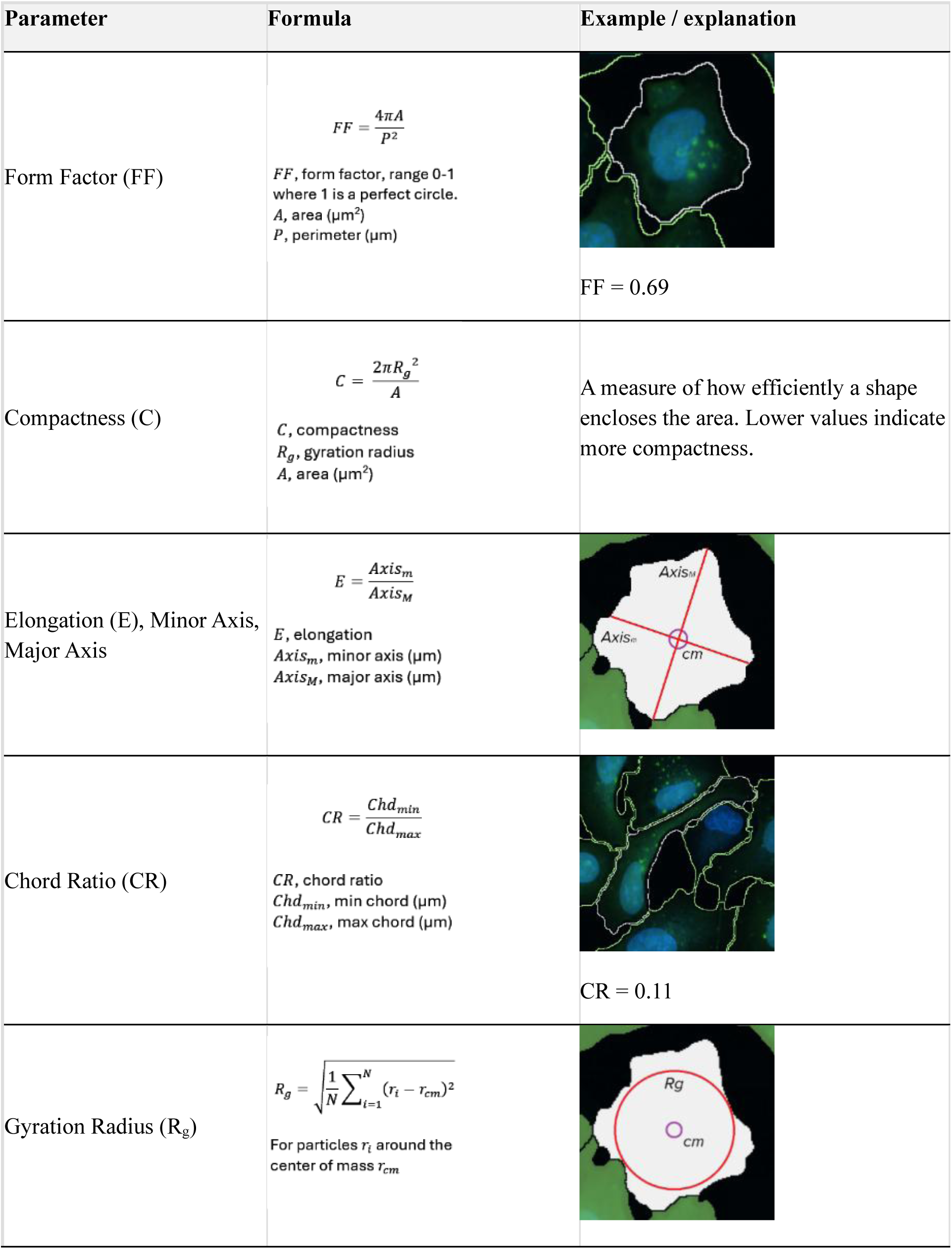
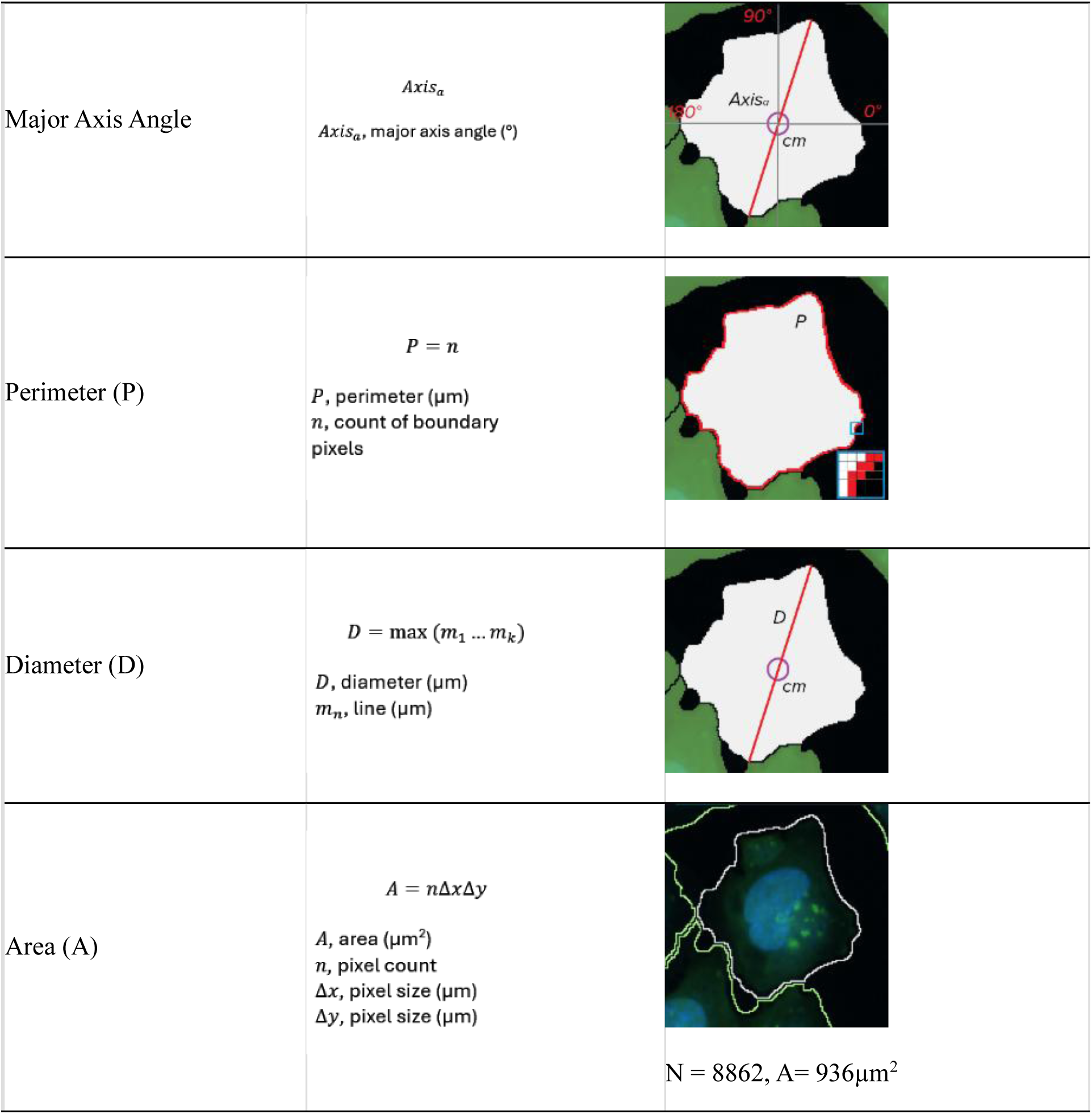
Image analysis parameters.

